# Structure-Activity Relationships, Tolerability and Efficacy of Microtubule-Active 1,2,4-Triazolo[1,5-*a*]pyrimidines as Potential Candidates to Treat Human African Trypanosomiasis

**DOI:** 10.1101/2023.03.11.532093

**Authors:** Ludovica Monti, Lawrence J. Liu, Carmine Varricchio, Bobby Lucero, Thibault Alle, Wenqian Yang, Ido Bem-Shalom, Michael Gilson, Kurt R. Brunden, Andrea Brancale, Conor R. Caffrey, Carlo Ballatore

## Abstract

Tubulin and microtubules (MTs) are potential protein targets to treat parasitic infections and our previous studies have shown that the triazolopyrimidine (TPD) class of MT-active compounds hold promise as antitrypanosomal agents. MT-targeting TPDs include structurally related but functionally diverse congeners that interact with mammalian tubulin at either one or two distinct interfacial binding sites; namely, the seventh and vinca sites, which are found within or between α,β-tubulin heterodimers, respectively. Evaluation of the activity of 123 TPD congeners against cultured *Trypanosoma brucei* enabled a robust quantitative structure-activity relationship (QSAR) model and the prioritization of two congeners for *in vivo* pharmacokinetics (PK), tolerability and efficacy studies. Treatment of *T. brucei*-infected mice with tolerable doses of TPDs **3** and **4** significantly decreased blood parasitemia within 24 h. Further, two once-weekly doses of **4** at 10 mg/kg significantly extended the survival of infected mice relative to infected animals treated with vehicle. Further optimization of dosing and/or the dosing schedule of these CNS-active TPDs may provide alternative treatments for human African trypanosomiasis.

## Introduction

Human African trypanosomiasis (HAT), also known as African sleeping sickness, is a life-threatening disease caused by the flagellated protozoan, *Trypanosoma brucei,* which is transmitted by tsetse flies. The two subspecies responsible for infection in humans are *T. b. gambiense* and *T. b. rhodesiense*. The disease is endemic in 36 sub-Saharan African countries with *T. b. gambiense* being the main infective agent in western and central regions, and *T. b. rhodesiense* in eastern and southern regions. Infection in humans progresses in two phases: an initial haemolymphatic stage (stage 1), and a meningoencephalitic stage (stage 2) when parasites cross the blood-brain barrier to infect the central nervous system (CNS). Stage 2 infection is nearly always fatal, if left untreated.^1–3^

Until recently, treatment of HAT has been limited to drugs with incomplete efficacy, severe toxicity, and difficult administration regimens.^4^ The recommendation of the nitroimidazole derivative, fexinidazole, by the European Medicines Agency^5^ in 2018 as the first-line oral treatment for both stages of HAT caused by *T. b. gambiense* was a major advance.^6, 7^ Also, recent encouraging clinical Phase II/III trial data with an investigational oxaborole drug, acoziborole, suggest that this compound may soon be registered as a new chemotherapeutic for combatting *T. b. gambiense* infection. ^8^ Nonetheless, given that trypanosomes have developed resistance to approved drugs in the past,^9–12^ the search for alternative treatments should remain a priority.

Tubulin and microtubules (MTs) are recognized as potential targets to treat parasitic infections, including trypanosomiases.^13–17^ The high degree of similarity between mammalian tubulin and the parasite counterpart (Figure 1) suggests that compounds that target mammalian MTs may also interfere with MT functions in the trypanosome. MT-targeting compounds form a broad category of biologically active compounds which include natural products and synthetic small molecules. These compounds exert potent anti-mitotic activity, especially in rapidly dividing cells, with often limited cell selectivity. Nevertheless, short-term treatments of cancer patients with MT-targeting compounds provide therapeutic benefits, as exemplified by the different MT-stabilizing and -destabilizing drugs that are employed in cancer chemotherapy.^18^ In contrast to the success of MT-targeting drugs in the treatment of different forms of cancer, data are sparse regarding the efficacy and tolerability of MT-active agents in the context of trypanosomal infections.^19^

**Figure 1.**
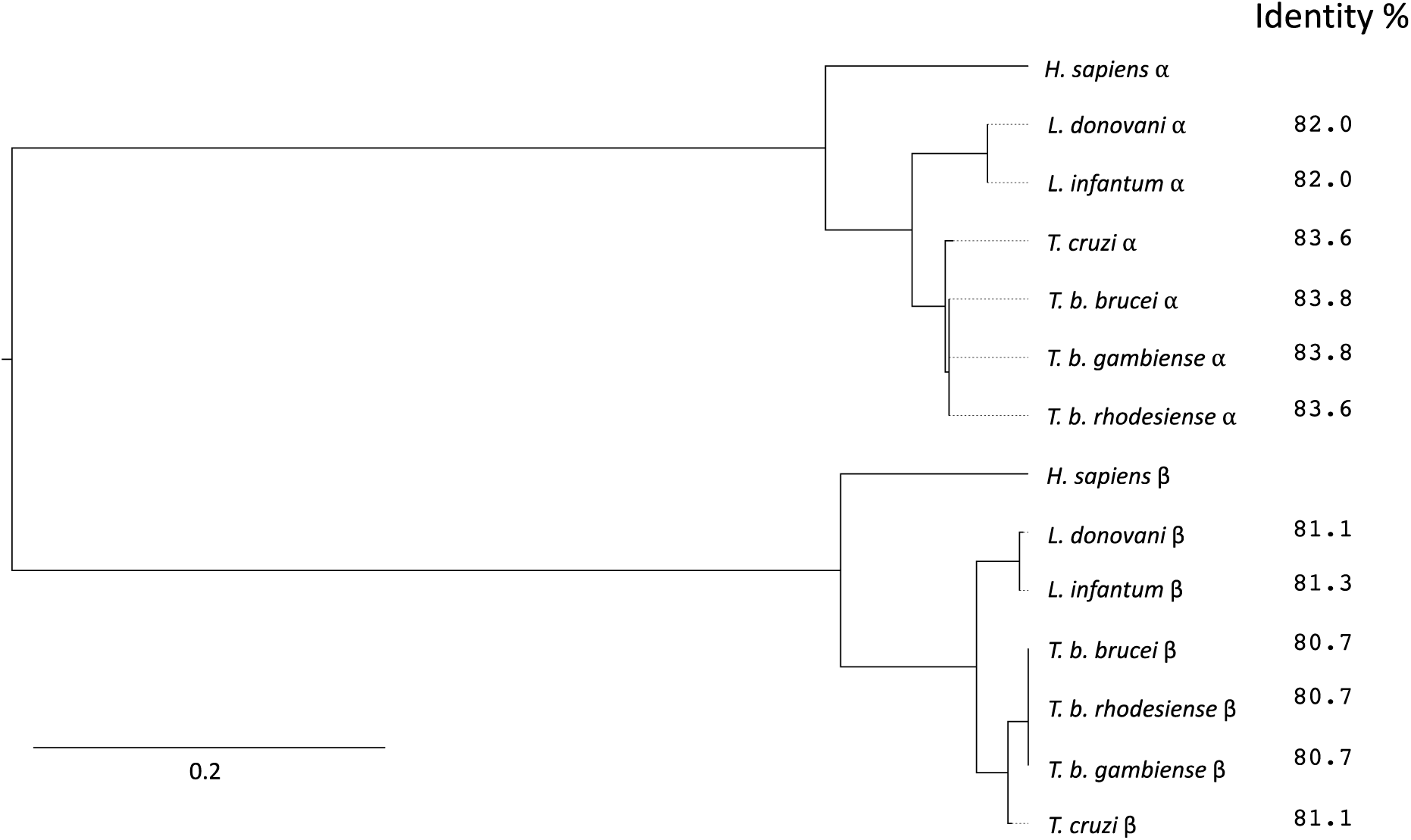
Phylogeny and sequence identity of α- and β-tubulin protein sequences of key trypanosomatid species. Maximum likelihood tree of α- and β-tubulin amino acid sequences from *T. b. brucei*, *T. b. gambiense*, *T. b. rhodesiense*, *Trypanosoma cruzi*, *Leishmania donovani*, *Leishmania infantum* and *Homo sapiens*. The software, MUSCLE^20, 21^, was used to align full-length tubulin sequences and served as the input for the construction of a maximum likelihood phylogenetic tree using IQ-TREE2.^22^ The interactive tree of life program, iTOL,^23^ was used to visualise the phylogenetic tree. The scale bar represents an approximate value for the amino acid substitution rate. Percentages indicate the identity of the α- and β-tubulin sequences of the different parasite species compared to their human orthologs, and were obtained using EMBOSS Water Pairwise Sequence Alignment.^21^

Our previous studies^24^ indicate that MT-active triazolopyrimidines (TPDs), a class of compounds that often have favourable ADME properties, including brain penetration,^25^ offer promise as potential anti-trypanosomal agents. In mammalian cells, these TPDs are believed to interact with either one or two distinct interfacial binding sites present within (the seventh site) or between (the vinca site) α,β-tubulin heterodimers to produce different effects on cellular tubulin and MTs. For example, studies with HEK-293 cells have shown that a subset of TPDs, referred to as Class I TPDs (*e.g.*, **1**, Figure 2), cause a dose-dependent increase in markers of stable MTs, such as acetylated α-tubulin (AcTub) and de-tyrosinated α-tubulin (GluTub), without altering total tubulin levels.^26^ X-ray co-crystal structure and computational studies have shown that Class I TPDs interact with the vinca site which is located between two longitudinally associated α,β-tubulin heterodimers.^27^ In contrast to Class I molecules, Class II TPDs (*e.g.*, **2** and **3**, Figure 2) induce a proteasome-dependent degradation of tubulin which, in HEK-293 cells, is often associated with a bell-shaped dose-response for both AcTub and GluTub.^28^ These compounds have a similar affinity for the vinca site as well as the seventh site^28^ (also known as the gatorbulin site^29^), which is an intradimer binding site between α- and β-tubulin subunits. Notably, recent studies have identified a third set of *hybrid* TPD congeners (*e.g.*, **4**, Figure 2), which, like Class II TPDs, are believed to interact with a similar affinity to both the vinca and seventh sites on tubulin, but like Class I TPDs, do not produce a significant decrease in total tubulin.^30^

**Figure 2.**
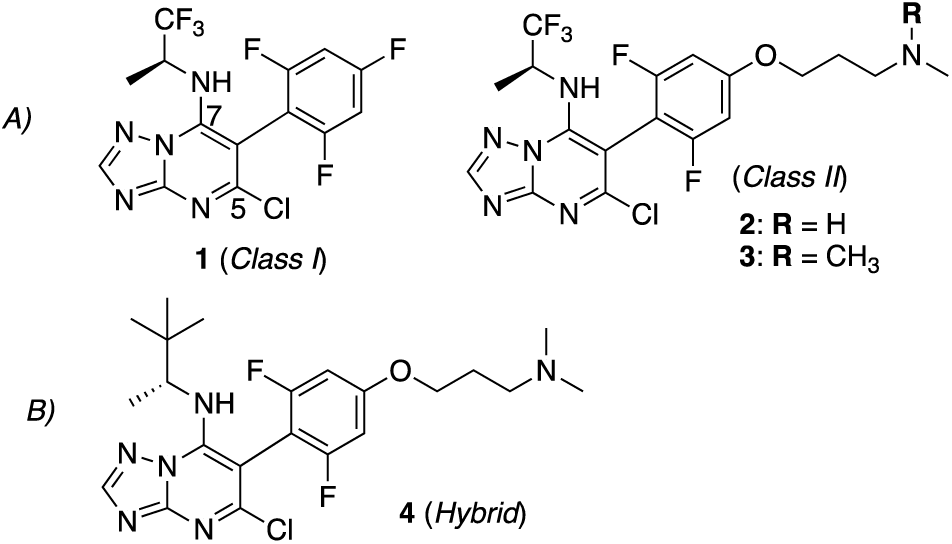
Structures of example MT-binding triazolopyrimidines. A) Class I and II TPDs that have been previously tested against *T. brucei in vitro*^24^; B) example of a hybrid TPD. Here, we report the activity of an expanded set of 123 TPDs, which includes several examples of Class I, Class II and hybrid compounds, against cultured *T. brucei*. This effort led to the development of a predictive QSAR model and the prioritization of selected Class II and hybrid TPD congeners for further assessment of PK, tolerability and efficacy in a mouse model of *T. brucei* infection. We show that one or two tolerable doses of appropriate TPDs decrease parasitemia and extend mouse survival. Furthermore, our data indicate that hybrid TPDs, which do not cause a reduction of total tubulin in mammalian cells, may provide a more favourable combination of efficacy and tolerability than Class II TPDs.

### Chemistry

1,2,4-Triazolo[1,5-a]pyrimidine analogs **1** – **123** (Table S1) were re-purposed from a drug discovery program for neurodegenerative diseases, and their synthesis has been described elsewhere.^25, 26, 30–33^

## Results

### Bioactivity of TPDs vs T. b. brucei Lister 427 and HEK-293 cells

The compound library evaluated in these studies (123 TPDs, Table S1) comprised 67 Class I, 36 Class II, and nine hybrid TPDs, as well as 11 congeners with no appreciable MT-stabilizing activity in HEK-293 cells. The *in vitro* killing activity of each compound against *T. b. brucei* Lister 427 was measured using the SYBR Green assay.^24^ Seventy-two compounds were active at or below the highest concentration tested of 4 μM. Of these, 22 compounds generated EC_50_ values of <250 nM (Table 1) with the HAT stage 1 drug, pentamidine, yielding an EC_50_ value of 69.64 ± 35.51 nM. The cytotoxicity of these top 22 compounds *vs*. HEK-293 cells was also evaluated (Table 1).

**Table 1.**
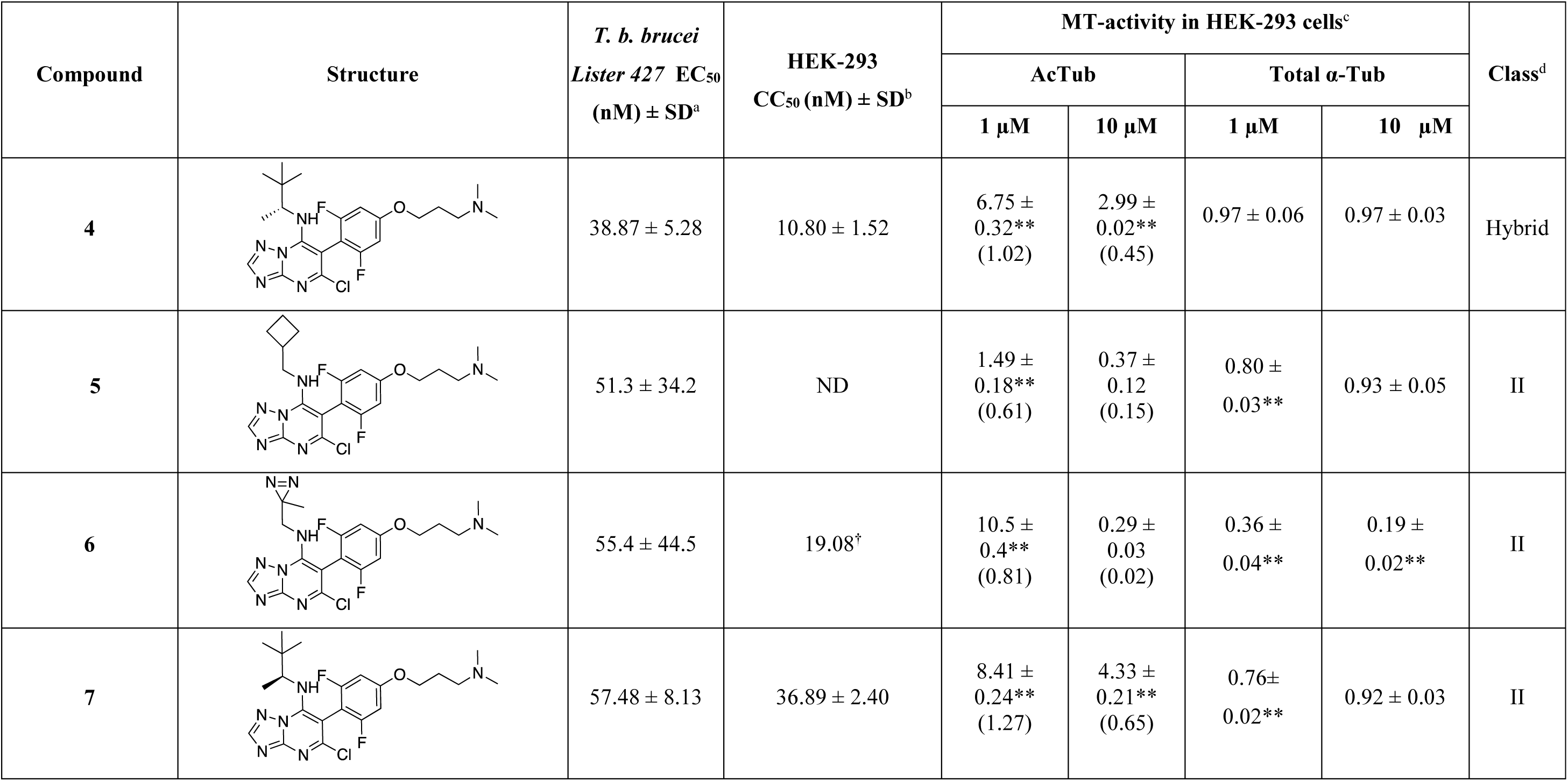
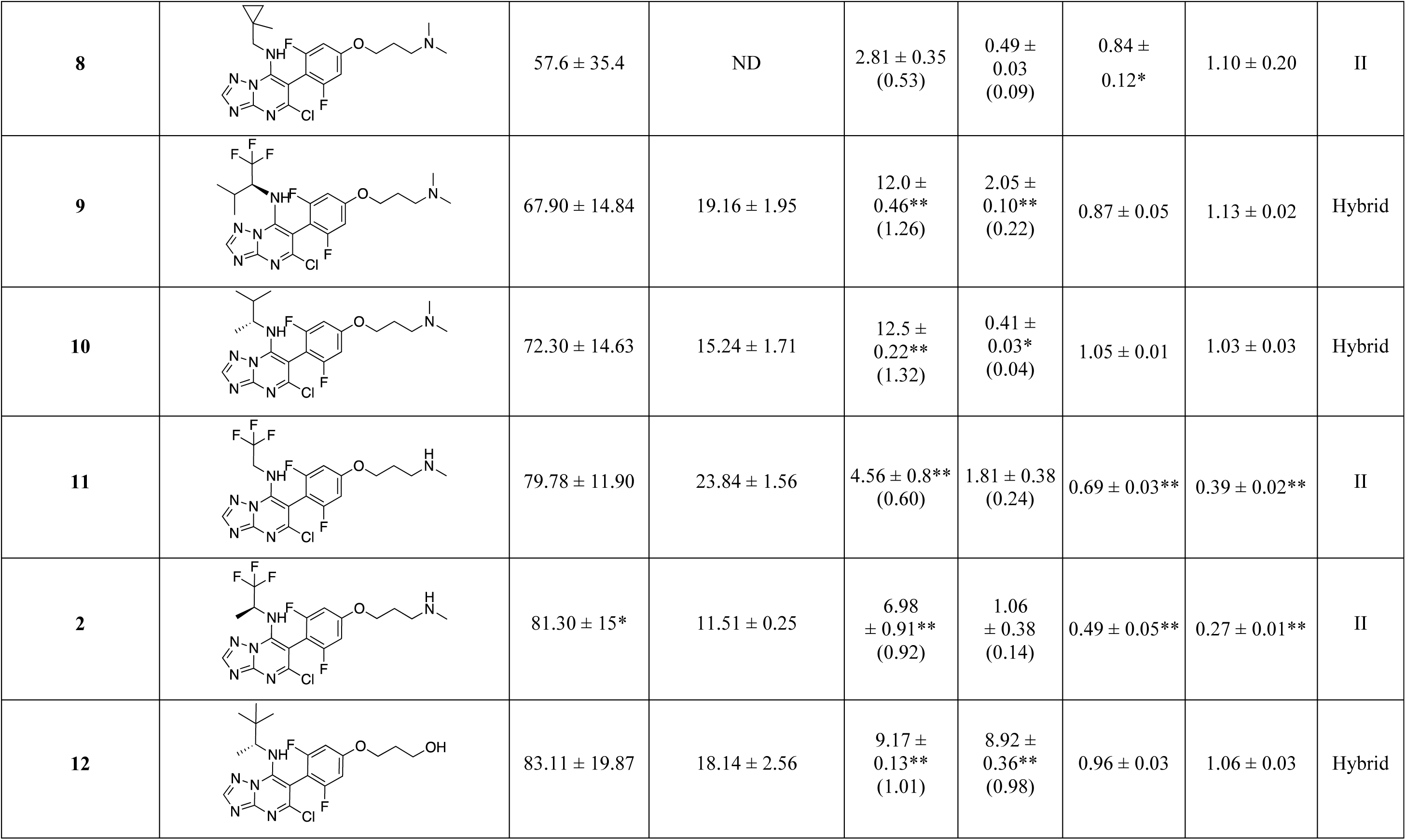
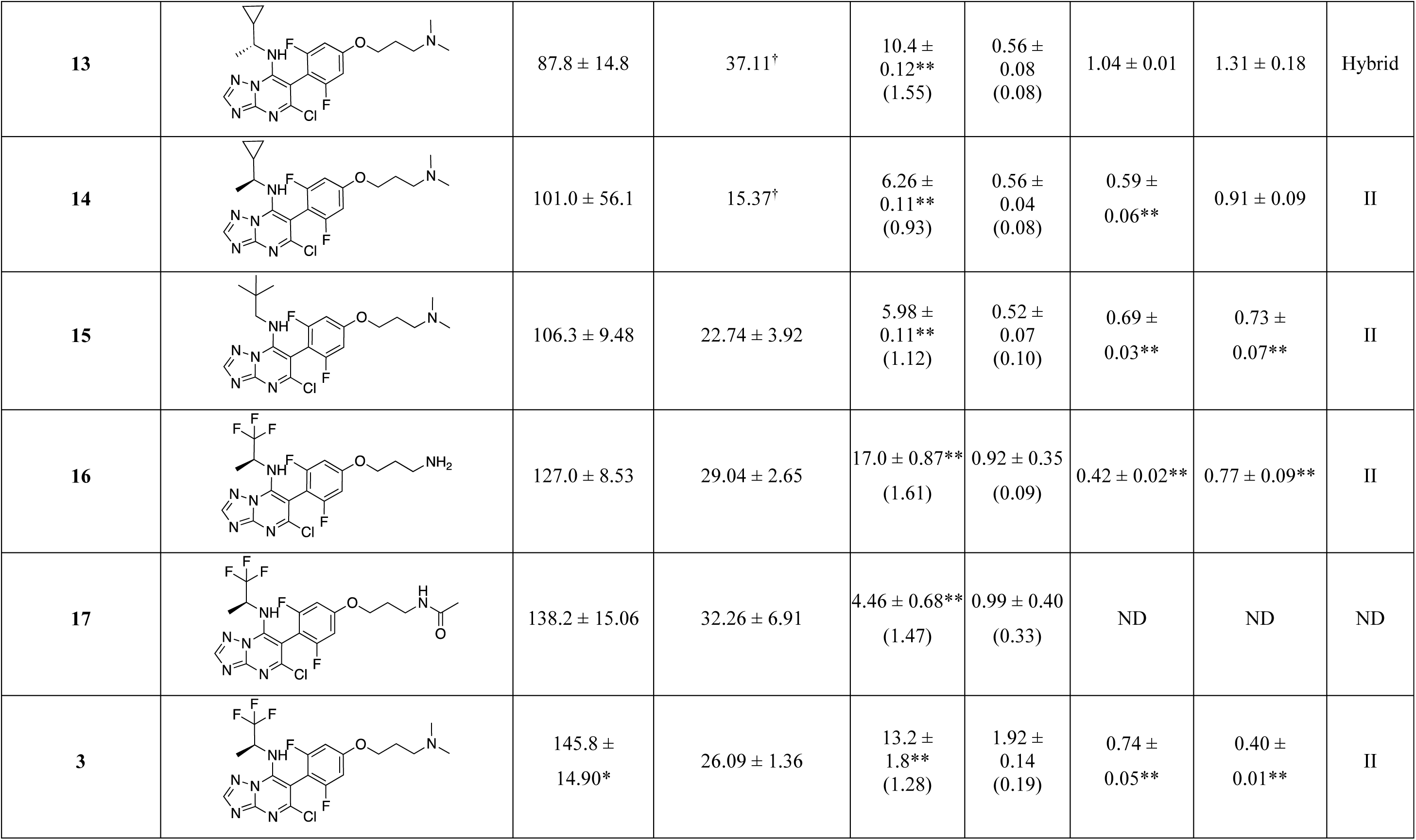
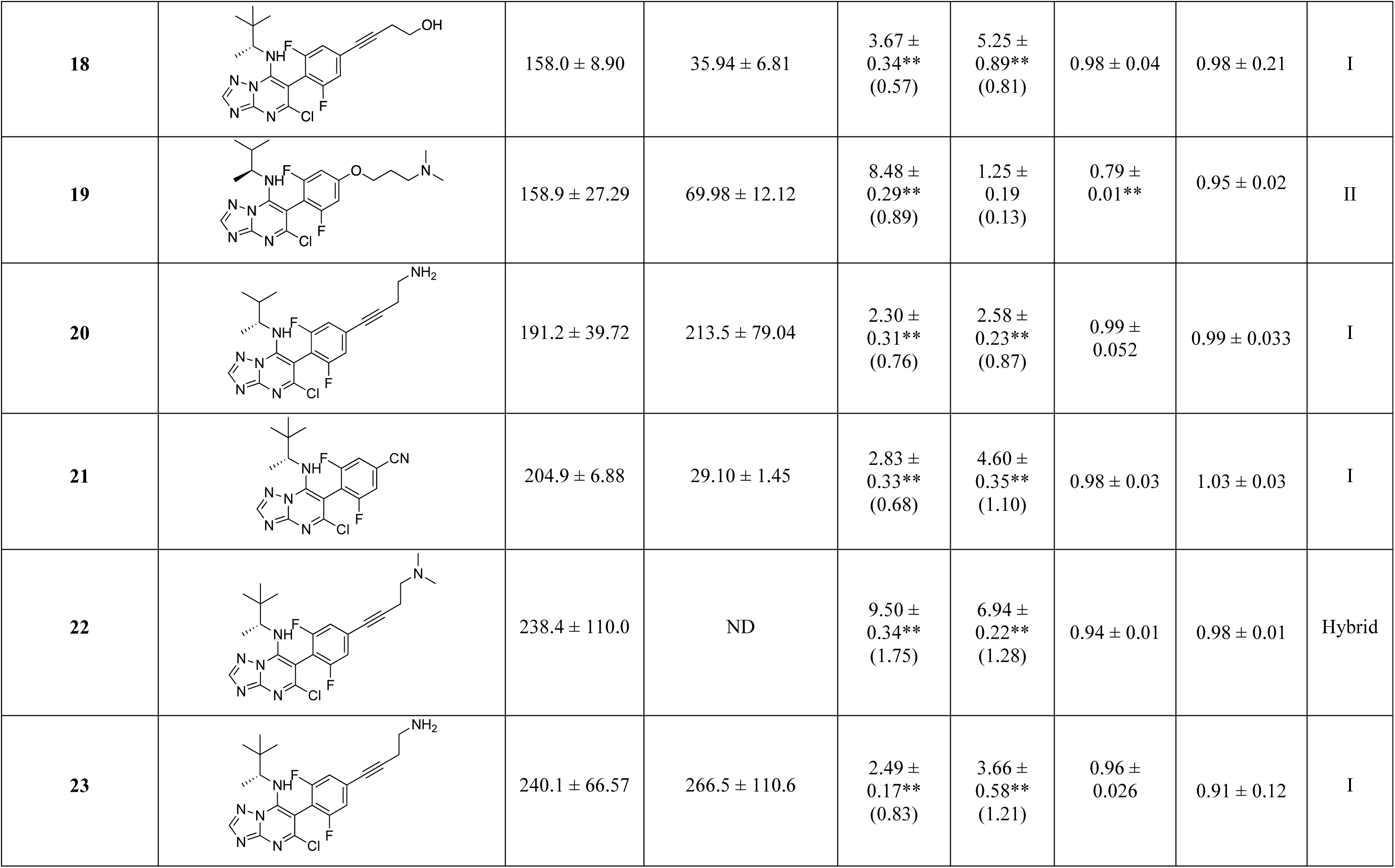
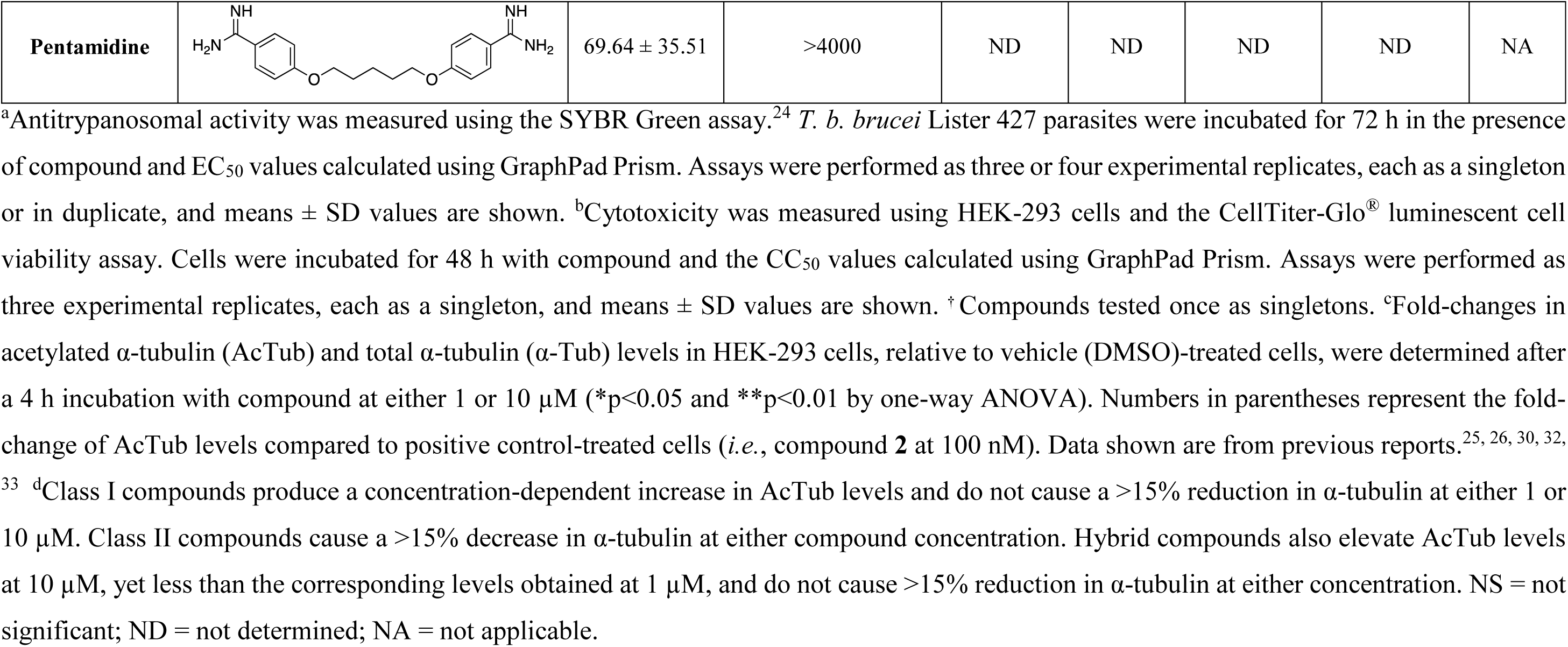
Antitrypanosomal activity, and cytotoxicity and MT-stabilizing activity in HEK-293 cells for selected TPD molecules (ranked based on their in vitro antitrypanosomal activity).

| Compound | Structure | *T. b. brucei*<br>*Lister* 427 EC<sub>50</sub><br>(nM) ± SD<sup>a</sup> | HEK-293<br>CC<sub>50</sub> (nM) ± SD<sup>b</sup> | MT-activity in HEK-293 cells<sup>c</sup> |  |  |  | Class<sup>d</sup> |
| --- | --- | --- | --- | --- | --- | --- | --- | --- |
|  |  |  |  | AcTub |  | Total α-Tub |  |  |
|  |  |  |  | 1 μM | 10 μM | 1 μM | 10 μM |  |
| 4 |  | 38.87 ± 5.28 | 10.80 ± 1.52 | 6.75 ± 0.32**<br>(1.02) | 2.99 ± 0.02**<br>(0.45) | 0.97 ± 0.06 | 0.97 ± 0.03 | Hybrid |
| 5 |  | 51.3 ± 34.2 | ND | 1.49 ± 0.18**<br>(0.61) | 0.37 ± 0.12<br>(0.15) | 0.80 ± 0.03** | 0.93 ± 0.05 | II |
| 6 |  | 55.4 ± 44.5 | 19.08<sup>†</sup> | 10.5 ± 0.4**<br>(0.81) | 0.29 ± 0.03<br>(0.02) | 0.36 ± 0.04** | 0.19 ± 0.02** | II |
| 7 |  | 57.48 ± 8.13 | 36.89 ± 2.40 | 8.41 ± 0.24**<br>(1.27) | 4.33 ± 0.21**<br>(0.65) | 0.76± 0.02** | 0.92 ± 0.03 | II |
| 8 | | $57.6 \pm 35.4$ | ND | $2.81 \pm 0.35$<br>(0.53) | $0.49 \pm 0.03$<br>(0.09) | $0.84 \pm 0.12^*$ | $1.10 \pm 0.20$ | II |
| 9 | | $67.90 \pm 14.84$ | $19.16 \pm 1.95$ | $12.0 \pm 0.46^{**}$<br>(1.26) | $2.05 \pm 0.10^{**}$<br>(0.22) | $0.87 \pm 0.05$ | $1.13 \pm 0.02$ | Hybrid |
| 10 | | $72.30 \pm 14.63$ | $15.24 \pm 1.71$ | $12.5 \pm 0.22^{**}$<br>(1.32) | $0.41 \pm 0.03^*$<br>(0.04) | $1.05 \pm 0.01$ | $1.03 \pm 0.03$ | Hybrid |
| 11 | | $79.78 \pm 11.90$ | $23.84 \pm 1.56$ | $4.56 \pm 0.8^{**}$<br>(0.60) | $1.81 \pm 0.38$<br>(0.24) | $0.69 \pm 0.03^{**}$ | $0.39 \pm 0.02^{**}$ | II |
| 2 | | $81.30 \pm 15^*$ | $11.51 \pm 0.25$ | $6.98 \pm 0.91^{**}$<br>(0.92) | $1.06 \pm 0.38$<br>(0.14) | $0.49 \pm 0.05^{**}$ | $0.27 \pm 0.01^{**}$ | II |
| 12 | | $83.11 \pm 19.87$ | $18.14 \pm 2.56$ | $9.17 \pm 0.13^{**}$<br>(1.01) | $8.92 \pm 0.36^{**}$<br>(0.98) | $0.96 \pm 0.03$ | $1.06 \pm 0.03$ | Hybrid |
| **13** | | $87.8 \pm 14.8$ | $37.11^{\dagger}$ | $10.4 \pm 0.12^{**}$<br>(1.55) | $0.56 \pm 0.08$<br>(0.08) | $1.04 \pm 0.01$ | $1.31 \pm 0.18$ | Hybrid |
| **14** | | $101.0 \pm 56.1$ | $15.37^{\dagger}$ | $6.26 \pm 0.11^{**}$<br>(0.93) | $0.56 \pm 0.04$<br>(0.08) | $0.59 \pm 0.06^{**}$ | $0.91 \pm 0.09$ | II |
| **15** | | $106.3 \pm 9.48$ | $22.74 \pm 3.92$ | $5.98 \pm 0.11^{**}$<br>(1.12) | $0.52 \pm 0.07$<br>(0.10) | $0.69 \pm 0.03^{**}$ | $0.73 \pm 0.07^{**}$ | II |
| **16** | | $127.0 \pm 8.53$ | $29.04 \pm 2.65$ | $17.0 \pm 0.87^{**}$<br>(1.61) | $0.92 \pm 0.35$<br>(0.09) | $0.42 \pm 0.02^{**}$ | $0.77 \pm 0.09^{**}$ | II |
| **17** | | $138.2 \pm 15.06$ | $32.26 \pm 6.91$ | $4.46 \pm 0.68^{**}$<br>(1.47) | $0.99 \pm 0.40$<br>(0.33) | ND | ND | ND |
| **3** | | $145.8 \pm 14.90^{*}$ | $26.09 \pm 1.36$ | $13.2 \pm 1.8^{**}$<br>(1.28) | $1.92 \pm 0.14$<br>(0.19) | $0.74 \pm 0.05^{**}$ | $0.40 \pm 0.01^{**}$ | II |
| 18 | | $158.0 \pm 8.90$ | $35.94 \pm 6.81$ | $3.67 \pm 0.34^{**}$<br>(0.57) | $5.25 \pm 0.89^{**}$<br>(0.81) | $0.98 \pm 0.04$ | $0.98 \pm 0.21$ | I |
| 19 | | $158.9 \pm 27.29$ | $69.98 \pm 12.12$ | $8.48 \pm 0.29^{**}$<br>(0.89) | $1.25 \pm 0.19$<br>(0.13) | $0.79 \pm 0.01^{**}$ | $0.95 \pm 0.02$ | II |
| 20 | | $191.2 \pm 39.72$ | $213.5 \pm 79.04$ | $2.30 \pm 0.31^{**}$<br>(0.76) | $2.58 \pm 0.23^{**}$<br>(0.87) | $0.99 \pm 0.052$ | $0.99 \pm 0.033$ | I |
| 21 | | $204.9 \pm 6.88$ | $29.10 \pm 1.45$ | $2.83 \pm 0.33^{**}$<br>(0.68) | $4.60 \pm 0.35^{**}$<br>(1.10) | $0.98 \pm 0.03$ | $1.03 \pm 0.03$ | I |
| 22 | | $238.4 \pm 110.0$ | ND | $9.50 \pm 0.34^{**}$<br>(1.75) | $6.94 \pm 0.22^{**}$<br>(1.28) | $0.94 \pm 0.01$ | $0.98 \pm 0.01$ | Hybrid |
| 23 | | $240.1 \pm 66.57$ | $266.5 \pm 110.6$ | $2.49 \pm 0.17^{**}$<br>(0.83) | $3.66 \pm 0.58^{**}$<br>(1.21) | $0.96 \pm 0.026$ | $0.91 \pm 0.12$ | I |
| Pentamidine |  | 69.64 ± 35.51 | >4000 | ND | ND | ND | ND | NA |

Consistent with our prior studies,^24^ Class I compounds were generally less potent against *T. brucei* compared to other TPD congeners. For the Class II TPDs and hybrid congeners tested, 18 compounds produced EC_50_ values of <1 µM with four Class II (**5**–**8**, Table 1) and three hybrid compounds (**4**, **9** and **10**, Table 1) exhibiting EC_50_ values comparable to or better than that for pentamidine. In contrast, none of the Class I TPDs was more potent than pentamidine in the *in vitro* assay, although several compounds (20 out of 67) yielded EC_50_ values <1 µM, including four (**18**, **20**, **21** and **23**, Table 1) with EC_50_ values in the 150–250 nM range.

### Quantitative Structure-Activity Relationships (QSAR)

To explore the relationship between the structural properties of the TPDs and their inhibition of the growth of *T. brucei*, we developed a QSAR model using a dataset of 91 congeners from the Class I, II and hybrid classes, and including examples of both active and inactive (*i.e.*, EC_50_ > 4 µM) compounds. The QSAR model was constructed using FieldTemplater (Flare, Cresset Software^34^) according to a field point template which describes the shape, electrostatic and hydrophobic properties of the molecules, and their spatial distribution.^35^ The field point template was generated by aligning and overlapping all molecules to maximise similarity in terms of electrostatic and molecular shape. The EC_50_ values generated in the *T. brucei* assay of the test compounds were expressed on a positive-logarithmic scale (pEC_50_) and correlated to field point distributions of the aligned molecules. After the alignment, the molecules were randomly partitioned into training and test sets comprising 63 and 28 compounds, respectively, and the QSAR model was validated by a “leave one out” (LOO) cross-validation process. The resulting QSAR model indicated a good predictive capability, as evidenced by both a robust regression coefficient (r^2^ = 0.96) and cross-validated coefficient of determination (q^2^ = 0.66) values with an r^2^ = 0.77 for the test set.

The QSAR model data provided useful information on the structural and electrostatic field points associated with antitrypanosomal activity (Figure 3). The substituent in the *para* position of the phenyl ring at C6 clearly had a strong influence on the biological activity as indicated by the number and the relatively large size of field points. In particular, the presence of the side chain at C6 was associated with an increase in activity (green field point), and the presence of a more positive electrostatic coefficient, such as stronger H-bond donors, appeared to further improve the antitrypanosomal activity of this family (cyan field point). Similarly, the presence of a favourable hydrophobic region in C7 (green field point) improved antitrypanosomal activity.

**Figure 3.**
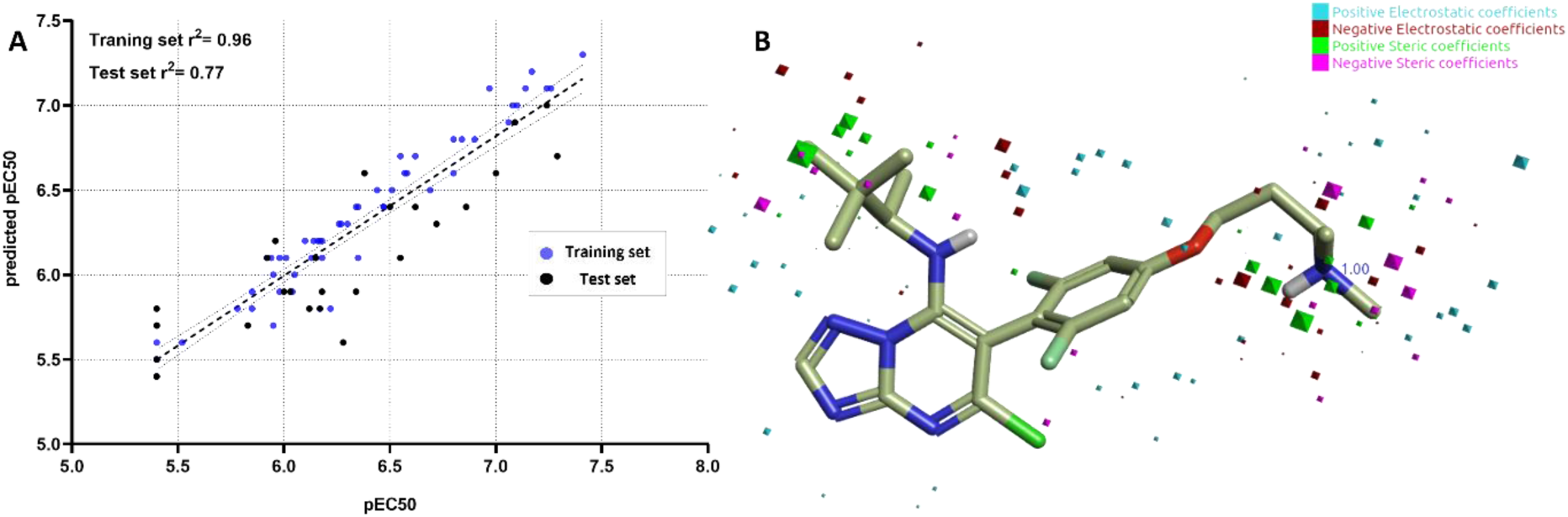
A) Experimental *vs.* predicted pEC_50_ values of the training and test sets of compounds derived from the QSAR model; B) The structure of TPD **4** with a visual representation of the QSAR’s electrostatic and steric coefficients which modulate the antitrypanosomal activity. The positive and negative electrostatic coefficients indicate whether a more negative or positive electrostatic potential field leads to an increase in activity, whereas the positive and negative steric coefficients indicate whether a greater steric bulk group leads to an increase or decrease in activity.

### Homology model of T. brucei tubulin

As an aid to the future design of ligands with potentially improved affinity for *T. brucei* tubulin, we investigated the binding mode of TPDs in the vinca and seventh site of *T. brucei*. Although there are no X-ray structures of *T. brucei* MTs, the *T. brucei* and human tubulin amino acid sequences share >80% identity (Figure 1), and, accordingly, we utilized a homology model of the vinca and seventh site of *T. brucei* constructed from a crystal structure of the Class II TPD, **2**, bound to a mammalian tubulin preparation (7CLD^28^). Overall, the *T. brucei* MT model exhibited a good degree of similarity with the mammalian template in the vinca and seventh sites due to a similar spatial distribution of several binding-site residues (Figure 4). However, some important differences were noted, especially in the vinca site. Thus, the Val353 residue of mammalian tubulin is replaced with Cys353 in *T. brucei*. Also, the Tyr222 and Ile332 residues are replaced in *T. brucei* with Phe222 and Val332, respectively. Combined, these amino acid changes in the trypanosome vinca site result in a different chemical reactivity (*cf.*, Val *vs.* Cys) and a significantly larger hydrophobic surface compared to the corresponding site in mammalian MTs. These distinguishing features may aid the optimization of TPD congeners with improved affinities for the trypanosome vinca site, *e.g.*, by altering the size/lipophilicity of TPD substituents and/or targeting the Cys353 residue.

**Figure 4.**
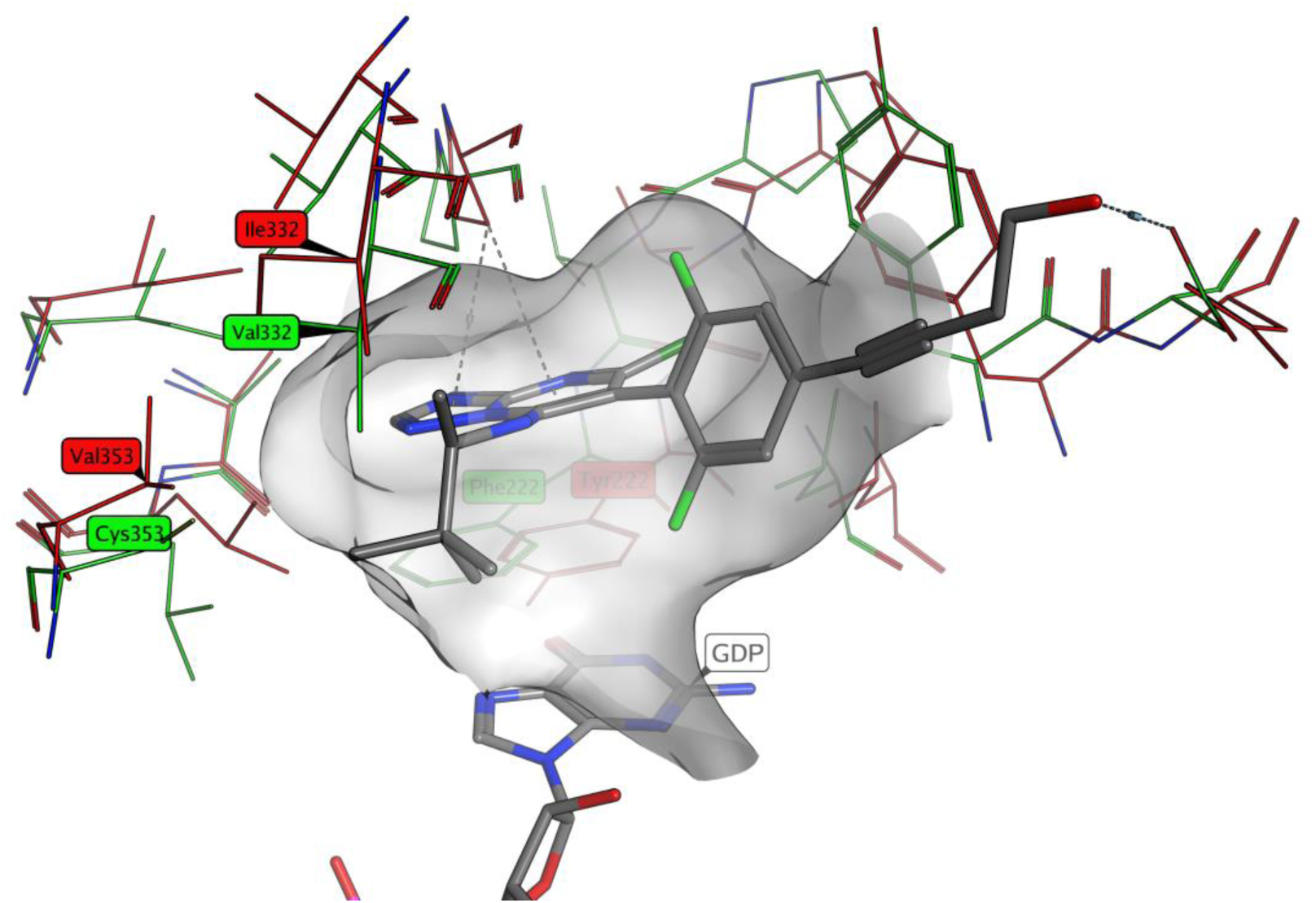
The docked structure of the Class I TPD, **18**, within the mammalian (red) and *T. brucei* (green) vinca site indicates that the Cys353 residue may be in proximity to the amine fragment at the C7 position of the TPD scaffold.

Based on the *in vitro* anti-trypanosomal activity data, a matched molecular pair of TPDs, namely, the Class II compound, **3**, and the hybrid compound, **4**, which differ structurally by the nature of the amine fragment at the C7 position of the TPD core, were selected for further studies; specifically (a) *in vitro* time-kill experiments, (b) brain and plasma pharmacokinetics (PK) assessments, (c) *in vivo* tolerability and (d) *in vivo* efficacy studies.

### Time-kill experiments

To investigate the speed of trypanocidal activity of the Class II TPD **3** and the hybrid TPD **4**, relative to the current drug, pentamidine, we measured parasite viability in the presence of a single and increasing concentrations of compounds as a function of time (3 – 72 h, Figure 5 and Figure S2). These side-by-side comparisons revealed that hybrid compound **4** was the fastest acting compound. For example, at the 12 h time point, parasite viability was ∼1.0 % when incubated with 4 µM compound **4**, compared to 23.9 and 52.8% in the presence of **3** and pentamidine, respectively (Figure 5A), and a similar trend was noted at 16 and 24 h. Further, after a 12 h incubation, the EC_50_ values for **3** and **4** were 6- and 13-fold less, respectively, than the value for pentamidine (Figure 5B).

**Figure 5.**
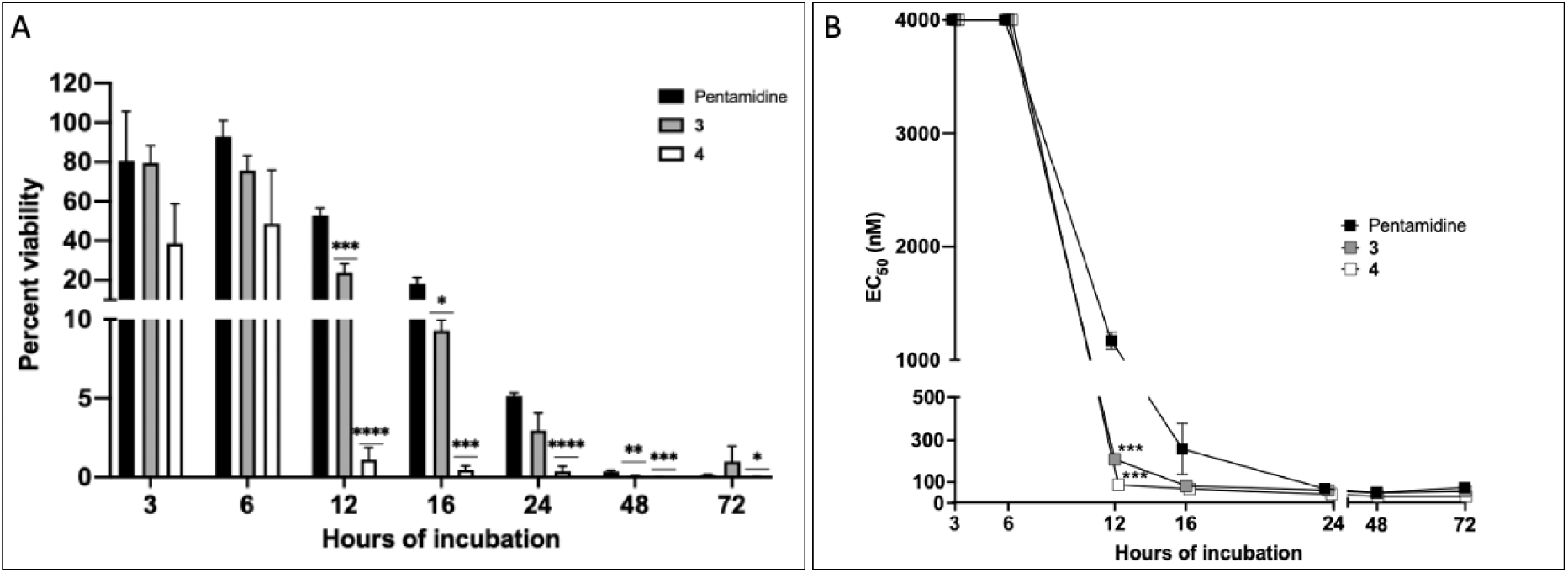
Time-kill assays for *T. b. brucei* Lister 427 in the presence of the TPDs **3** and **4**, and pentamidine. (**A**) Percentage viability was measured in the presence of 4 µM compound. (**B**) *T. brucei* was exposed to 0.87 nM through 4 µM compound as a function of time. All assays were performed as three experimental replicates, each as a duplicate. The activity of test compounds was normalized to DMSO controls from the same plate and means ± SEM values are shown. The Student’s unpaired *t*-test was used to compare the data for the TPDs *vs.* those for pentamidine: *p < 0.05, **p < 0.01, ***p < 0.001, ****p < 0.0001.

### PK studies

Hybrid compound **4** was evaluated for brain and plasma PK after a 2 mg/kg intraperitoneal (i.p.) injection (Figure 6). These studies revealed a brain-to-plasma AUC value of 0.63. Compound **3** had a lower brain-to-plasma AUC value of 0.37, as previously reported.^25^

**Figure 6.**
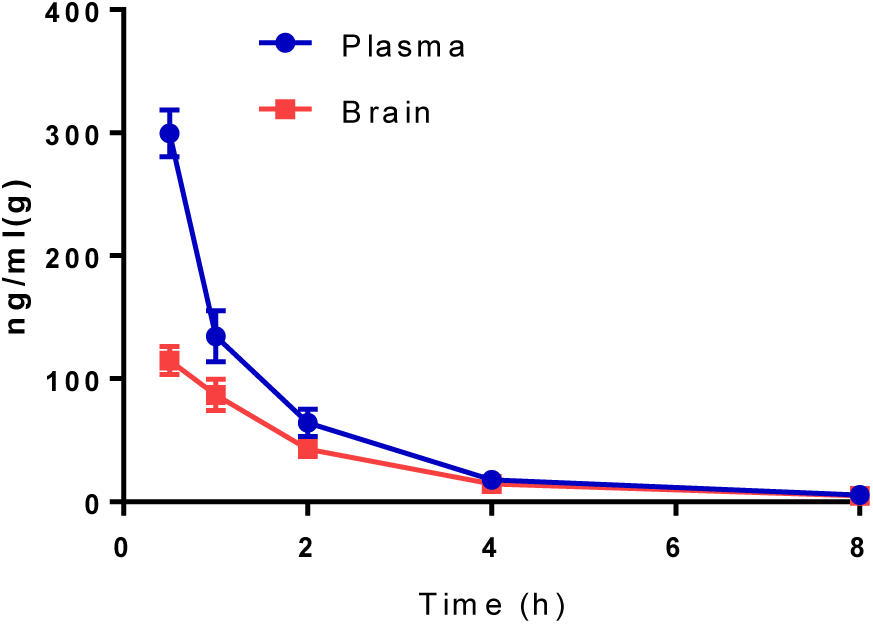
Brain and plasma PK of **4** after an i.p. administration of 2 mg/kg. Data are from one experiment with three mice per time point.

### Tolerability studies

To identify tolerable doses of TPDs in advance of efficacy studies, an initial dose range-finding experiment was conducted that involved the i.p. administration of single and increasing doses of **3** and **4** to BALB/c mice (*n =* 1). Animals were monitored twice daily for up to 48 h post-treatment for signs of intolerance. The highest doses of **3** and **4** not producing overt signs of intolerance ranged from 10 to 15 mg/kg, and from 15 to 20 mg/kg, respectively. Follow-up experiments were then conducted with groups of six female and male mice per compound. Compounds **3** and **4** were administered at doses ranging from 6.6 to 12.2 mg/kg and from 8.9 to 18.9 mg/kg, respectively. Regardless of sex, doses exceeding 10 and 17 mg/kg for **3** and **4,** respectively, induced signs of intolerance (reduced grooming and socialization) over the 48 h surveillance period and the mice were humanely euthanized.

### In vivo efficacy studies

Six-week-old BALB/c female mice were infected i.p. with 1 × 10^5^ parasites. Once parasitemia was established by day 2 post-infection (approximately 1–5 × 10^7^ parasites/ml of blood; Figure 7), mice were divided into groups of five, and each group received a single i.p. dose of compound at the concentrations described below. Infected control mice groups were those that received either no treatment or given vehicle only (vehicle controls). For both sets of controls, parasitemia had increased to between 1 × 10^8^ and 1 × 10^9^ parasites/ml of blood by day 3 post-infection (Figure 7 and Table S2) and were humanely euthanized as, otherwise, they would have died within 24 h. Pentamidine, at 4 mg/kg, decreased parasitemia to below the detectable limit (*i.e.,* 2.5 × 10^5^ parasites/ml) within 24 h of administration (Figure 7A). TPD **3**, at 5, 7.5 and 10 mg/kg, significantly decreased parasitemia levels relative to the infected untreated control after 24 h, and, for the two higher doses, parasitemia was below the detection limit (Figure 7B). TPD **4**, was also administered at 5, 7.5 and 10 mg/kg (Figure 7C). The compound was ineffective at the lowest dose but produced a significant reduction in parasitemia at the two higher doses, with the 10 mg/kg dose resulting in parasitemia being below the detection limit.

**Figure 7.**
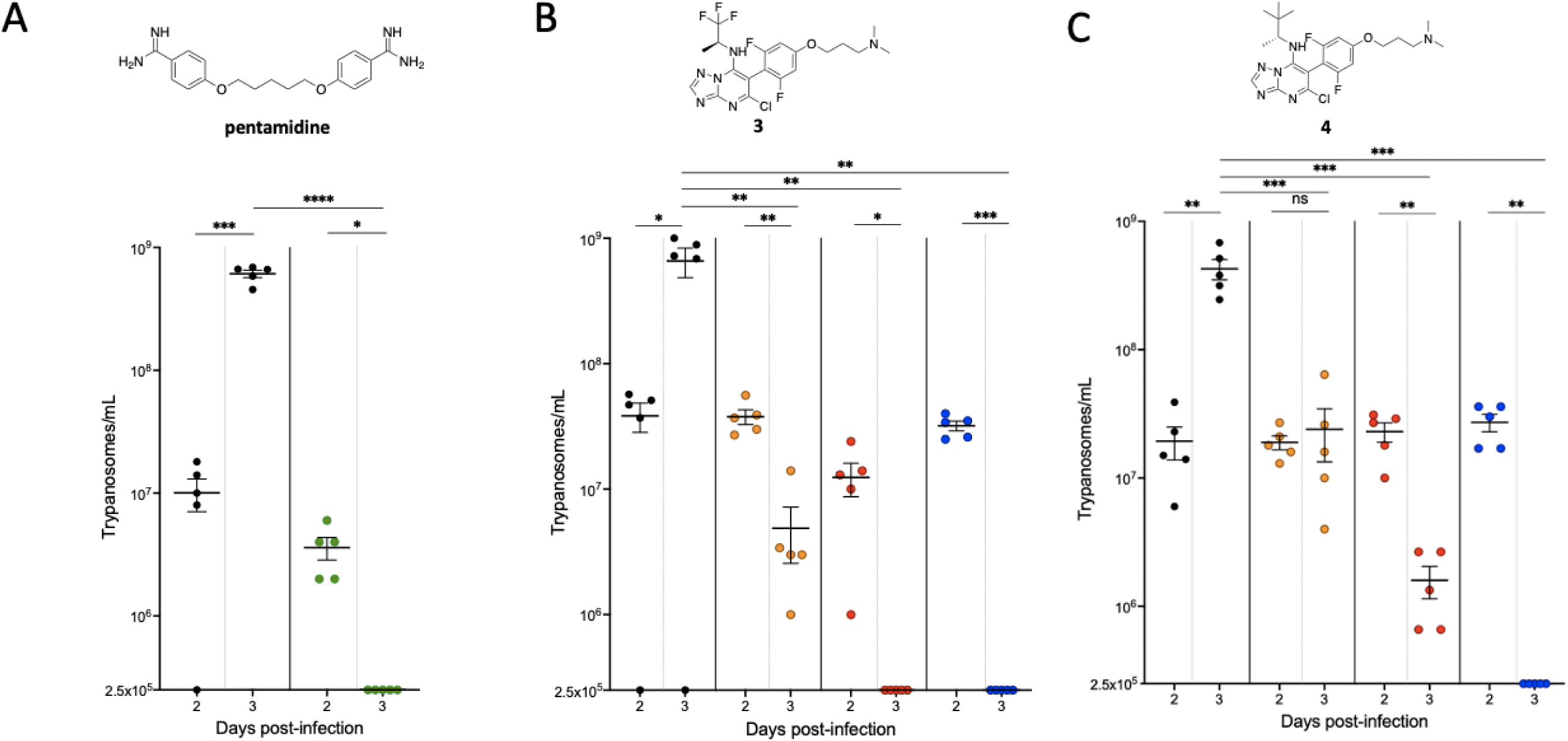
Effect of TPD treatment on blood parasitemia in mice infected with *T. brucei.* Female BALB/C mice were infected i.p. with 1×10^5^ *T. b. brucei* Lister 427 parasites. On day 2 post-infection, mice were divided into groups of five and treated with an i.p. injection of (A) 4 mg/kg pentamidine, or 5 mg/kg (orange), 7.5 mg/kg (red) or 10 mg/kg (blue) of **3** (B) or **4** (C). Black points indicate the corresponding infected vehicle controls. Parasitemia was measured as described in the experimental section and the limit of accurate detection was 2.5×10^5^ parasites/ml of blood. Data are expressed as means ± SEM. The Student’s paired *t*-test was used for intra-group testing for days 2 and 3 post-infection, and the Student’s unpaired *t*-test was used for inter-group testing on day 3 post-infection compared to the respective controls: *p < 0.05, **p < 0.01, ***p < 0.001, ****p < 0.0001, ns = not significant.

The survival time of infected mice post-treatment with compounds **3** and **4** at 5, 7.5 and 10 mg/kg was also measured to evaluate compound efficacy (Figure 8). All vehicle control animals were humanely euthanized when parasitemia exceeded 1 × 10^8^ parasites/ml. Pentamidine at 4 mg/kg was apparently curative as the mice survived to the end of the study (day 15 post-infection) without parasite recrudescence (Figure 8A and Table S3). For both TPDs **3** and **4** at the lowest and two lowest doses, respectively (Figure 8B and 8C), mouse survival was increased marginally to a maximum of day 7 post-infection. All mice that received the highest 10 mg/kg dose of each compound were still alive by day 9 post-infection, and, due to recrudescence of parasitemia (Tables S4 and S5), were administered a second dose. Specifically, TPD **3** was administered at 7 and 5 mg/kg to the 7.5 and 10 mg/kg groups, respectively, whereas in the case of TPD **4**, which appeared to be more tolerable at the higher doses compared to **3**, the 10 mg/kg group received an additional 10 mg/kg dose. TPD **3** extended the survival of 3/5 and 2/5 mice from the initial 7.5 and 10 mg/kg groups, respectively, to day 15 post-infection (Figure 8B), at which time blood parasitemia was again measurable and the mice were euthanized (Table S4). In contrast, all of the mice treated with the second dose of TPD **4** survived to day 16 post-infection (Figure 8C) and were euthanized due to parasite recrudescence in 4/5 mice (Table S5). These data demonstrate that both TPDs **3** and **4** provide a therapeutic benefit, with the latter being better tolerated at higher doses, and extending the survival time of the entire group of mice to four times that of the vehicle control group.

**Figure 8.**
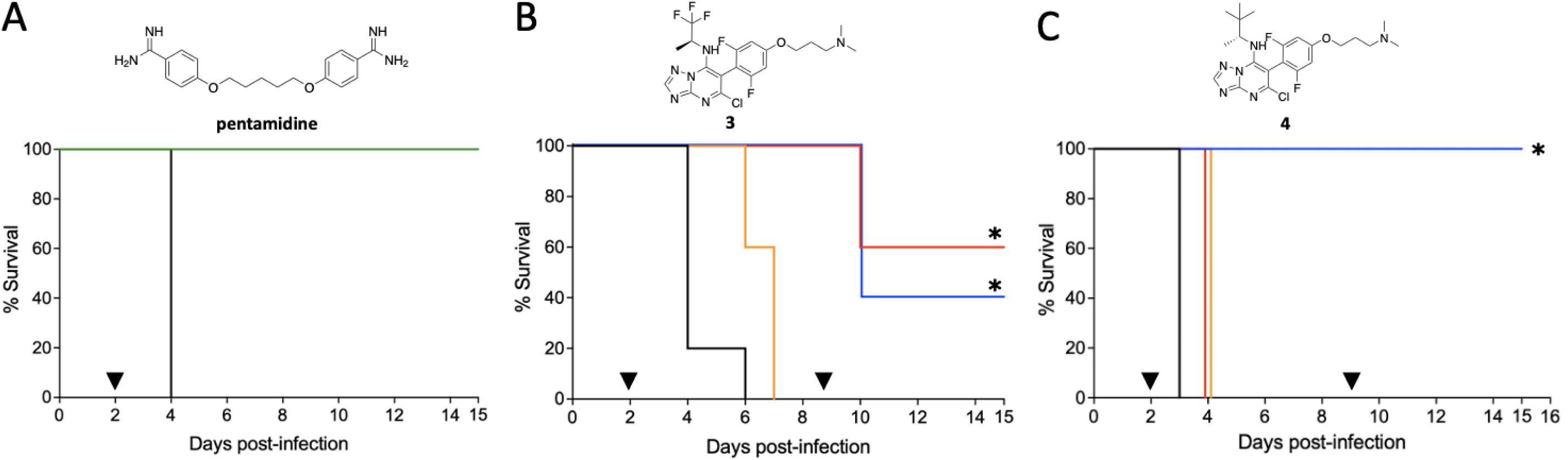
Kaplan–Meier curves for survival of *T. brucei*-infected mice treated with TPDs. Female BALB/C mice were infected i.p. with 1 × 10^5^ *T. b. brucei* Lister 427 parasites. On day 2 post-infection, when parasitemia was established, mice were divided into groups of five and treated with an i.p. injection of (A) 4 mg/kg pentamidine (green), or 5 mg/kg (orange), 7.5 mg/kg (red) or 10 mg/kg (blue) of **3** (B) or **4** (C). Black lines in each panel indicate the survival of infected mice treated with vehicle alone. An additional dose of TPD **3** or **4**, as indicated in the text, was administered on day 9 post-infection (▼) when parasitemia was detected. Asterisks (*) indicate the presence of blood parasitemia at the end of the study (days 15 and 16 post-infection for **3** and **4**, respectively; see Tables S3–S5).

## Discussion

Short-term treatments with MT-targeting compounds offer significant therapeutic value in cancer chemotherapy, despite the very limited cell selectivity of these anti-mitotic agents. Although MT-targeting compounds have shown potent *in vitro* antitrypanosomal activity, thus far, few studies^19^ have investigated the *in vivo* efficacy and safety of short-treatments with compounds of this type in the context of trypanosomal infections. Among the different classes of MT-targeting compounds, TPDs are attractive due to their generally favourable drug-like properties, including oral bioavailability and brain penetration. In recent years, TPDs have been the focus of intense studies as candidate therapeutics for neurodegenerative diseases.^25, 26, 33^ Moreover, recent mechanism of action^27, 28^ and SAR^26, 33^ studies have revealed that these molecules can produce significantly different effects on tubulin/MT structures in mammalian cells, due to their ability to interact with either one or two distinct interfacial binding sites: the vinca site and the seventh site.^29^ Indeed, depending on the choice of substituents around the TPD core, these compounds show different affinities for the two binding sites. For example, the TPDs that are expected to show greater affinity for the vinca site generally promote tubulin polymerization in cells leading to the formation of relatively stable MTs in a dose-dependent manner. Unlike these Class I TPDs, other structurally related congeners show little or no preferential binding for either the vinca or the seventh sites. In cell-based studies of MT-stabilization, these compounds are generally characterized by an unusual bell-shaped dose response relationship with respect to cellular markers of stable MTs, such as AcTub and GluTub. Among these dual vinca/seventh site binders, the Class II TPDs produce a proteasomal-dependent degradation of total tubulin, whereas other, more recently identified hybrid TPDs do not alter total tubulin levels.^30^

Our previous *in vitro* antitrypanosomal studies^24^ with representative examples of Class I (**1**) and Class II (**2** and **3**) TPDs have shown that the latter TPDs can reach relatively potent antiparasitic activities with EC_50_ values in the sub-µM range. The studies presented herein focused on evaluating the *in vitro* antitrypanosomal activity of an expanded set of 123 compounds, which include examples of the new hybrid TPDs.^30^ This screening effort facilitated a predictive QSAR model and identified several TPDs with potent antitrypanosomal activity (*i.e.*, EC_50_ <250 nM, Table 1). In general, the Class I TPDs appeared to be comparatively less active than either the Class II or hybrid TPDs. Among the seven TPD analogues with *in vitro* EC_50_ values either comparable to or better than that for pentamidine, four were Class II and three were hybrids (Table 1).

To investigate whether short-term treatments with Class II and hybrid TPDs can produce therapeutic benefits in a mouse model of *T. brucei* infection, a matched molecular pair of compounds, **3** and **4**, was evaluated. Subsequent to establishing the tolerable doses, treatment with these compounds resulted in a dose-dependent decrease in parasitemia, including to below detectable levels, and extended the survival of infected mice relative to vehicle controls. In the case of hybrid compound, **4**, two once-weekly doses of 10 mg/kg resulted in an extension of survival to day 16 post-infection when the study concluded. Overall, the efficacy data suggest that despite the absence of *in vitro* cell selectivity index (as evidenced in Table 1), short-term treatments with TPDs offer therapeutic value against trypanosomiases. In this context, our results suggest that hybrid TPDs may be more promising than Class II TPDs in terms of their tolerability and efficacy, and that further optimization of these compounds is warranted. Finally, although our studies have so far focused on the evaluation of the efficacy and tolerability of short-term treatments with MT-targeting TPDs that have not been synthetically optimized to engage trypanosome tubulin, structural analysis via homology modelling has highlighted significant differences in the structure and characteristics of the vinca site between the trypanosomal and mammalian MTs that could be exploited for the design of congeners with increased selectivity for the parasitic tubulin.

### Conclusions

We evaluated the *in vitro* antitrypanosomal activity of 123 1,2,4-triazolo[1,5- *a*]pyrimidine analogues with MT-stabilizing activity. The *in vitro* screening effort identified seven compounds with EC_50_ values against *T. brucei* that were comparable to, or better than pentamidine, the drug for stage 1 HAT. A matched molecular pair of TPDs was selected for further side-by-side evaluation of *in vivo* PK, tolerability and efficacy. Doses of 7.5 and 10 mg/kg of the Class II TPD, **3**, or 10 mg/kg of the hybrid TPD, **4**, in mice infected with *T. brucei* decreased parasitemia to levels below the limit of detection within 24 h. Notably, administration of **4** at 10 mg/kg once a week for two weeks to mice extended survival without overt signs of toxicity. Taken together, these results have (*a*) established an *in vitro* - *in vivo* activity correlation for the antitrypanosomal activity of TPDs, (*b*) demonstrated that one or two tolerable doses of an appropriate TPD results in significant antitrypanosomal effects *in vivo*, and (*c*) suggest that MT-targeting TPDs that do not cause a proteasome-dependent reduction of total tubulin may provide a more favourable combination of efficacy and tolerability. Finally, the data generated in these studies facilitated the development of a robust QSAR model that could be used for the further exploration of additional antitrypanosomal TPD congeners.

## Experimental section

### Materials and Methods

Compounds **1**–**123** were synthesized previously in our laboratories as part of a drug discovery program for Alzheimer’s disease.^25, 26, 30, 32, 33^

### Computational Studies

The 3D structures of TPDs were prepared considering their ionization states at pH 7 ± 2 and imported into the Flare software (Cresset Inc., Cambridgeshire, UK)^36^ for setting the field-based 3D-QSAR model. Each compound was aligned to a pharmacophore model generated using the FieldTemplater module of Flare.^37^ The pharmacophore model was determined by comparing the electrostatic and hydrophobic properties of the most active compounds. The alignments were performed to maximize similarity in terms of molecular field and molecular shape similarity. After the alignment, the molecules were randomly partitioned between the Training set (61) and test set (28), using the stratified sampling of activity values. The biological activity was expressed as p= - log(EC_50_). The 3D-QSAR was constructed based on the aligned molecules, and it was assessed by the LOO technique to optimize the activity prediction model.^38^

The homology model of *T. brucei* tubulin was generated using the Molecular Operating Environment (MOE2022.02)^39^ homology model tool suite. The FASTA sequences of the α- (P04106) and β- (P04107) tubulins were downloaded from the UniProt^40^ database and modelled on the template of the mammalian tubulin crystal structure with co-crystallized ligands (PDB: 7CLD). The final homology model was energy-minimized to a root mean square (RMS) gradient of 0.01 kcal/mol/Å2 using the AMBER10:EHT force field. The model was verified by the Structural Analysis and Verification Server (SAVES v6.0)^41^ to evaluate its stereo-chemical quality (Figure S3). The docking studies of the TPDs were performed using the predicted protein structure. Prediction of the protonation states of protein residues was calculated using PrepWizard (Schrödinger 2022.2)^42^ considering a temperature of 300 K and a pH of 7. An Å docking grid (inner-box 10Å and outer-box 20 Å) was prepared using as centroid the co-crystallized ligands. The docking studies were performed using Glide SP precision (Schrödinger 2022.2)^43^ keeping the default parameters and settings. MOE 2022.02 was used to visualize the structures and acquire the images, and Prism GraphPad^44^ and Data Warrior^45^ were used for the statistical analysis.

### Maintenance of *In Vitro* Cell Cultures

#### T. brucei

Bloodstream forms of *T. b. brucei* Lister 427 were grown in 5 ml of modified HMI-9 medium^46^ in T25 suspension cell culture flasks (Genesee Scientific, Cat. No. 25-213) under a humidified atmosphere of 5% CO_2_ at 37 °C. Parasites were maintained in log-phase growth (between 1×10^5^ and 1×10^6^ parasites/ml) and sub-cultured every 48 h.^47^

#### HEK-293

These were cultured in Dulbecco’s Modified Eagle’s Medium (DMEM, Gibco, Cat. No. 11965092) containing 10% heat-inactivated fetal bovine serum (Invitrogen, Cat. No. 16140-071) and 1% penicillin-streptomycin (Gibco, Cat. No. 15140122) at 37 °C and 5% CO_2_. Cells were grown in T75 cell culture flasks (Thermo Scientific, Cat. No. 156499) and sub-cultured when 70–80% confluent.

### *In Vitro* Assay Protocols

*SYBR Green Viability Assay With T. brucei.* Growth inhibition of *T. b. brucei* Lister 427 after compound exposure was determined using established procedures.^24, 47^ The test compounds were serially diluted in DMSO (Thermo Scientific™, Cat. No. 85190) across eight concentrations ranging from 4 μM to 0.87 nM. Trypanosomes in log-phase growth were suspended at 2×10^5^ parasites/ml in HMI-9 medium under continuous agitation. Cells were dispensed as 100 μl aliquots into 96-well black or white plates with flat and clear bottoms (Corning^®^, Cat. No. 3903) containing 1 μl of test compound in 0.5% DMSO. An additional 100 μl medium was added, and the plates were then incubated for 72 h at 37 °C and 5% CO_2_. Trypanosomes were lysed by the addition of 50 μl/well of lysis solution (30 mM Tris pH 7.5, 7.5 mM EDTA, 0.012% saponin (Alfa Aesar, Cat. No. A18820), and 0.12% Triton X-100 (Sigma Aldrich, Cat. No. T8787) containing 0.3 μl/ml SYBR Green I (10,000× in DMSO; Lonza, Cat. No. 50513). Plates were incubated in the dark for at least 1 h at room temperature. Fluorescence was measured using a 2104 EnVision^®^ multilabel plate reader (PerkinElmer, Waltham, MA) or a Synergy HTX multi-mode reader (485/535 nm, excitation/emission wavelengths). The activity of test compounds was normalized to those of the DMSO controls from the same plate. Dose–response curves and EC_50_ values, *i.e.,* the concentration of compound required to generate a 50 % inhibition of the parasite growth rate, were calculated using GraphPad Prism version 9.4.1 for MAC OS (GraphPad Software, San Diego, CA, www.graphpad.com). Assays were performed as three or four experimental replicates, each as a singleton or in duplicate.

#### HEK-293 Counter-Screen Aassay

The inhibition of cell proliferation was determined using the CellTiter-Glo^®^ reagent (Promega, Cat. No. G7572) according to the manufacturer’s recommendation. Compounds in 100% DMSO were diluted in 50 μL DMEM medium using 96-well white plates with flat and clear bottoms (Corning^®^, Cat. No. 3903) and a further 50 μL of cells in DMEM were added such that the final DMSO concentration was 1%. Eight-point concentration-response assays (depending on the initial EC_50_ value, ranging from 4 μM to 0.87 nM) were set up. HEK-293 cells were then diluted to 1×10^5^ cells/ml in DMEM and dispensed into the previously prepared 96-well plates at 50 μL per well. After 48 h at 37 °C and 5% CO_2_, cells were lysed by the addition of 50 μL/well of CellTiter-Glo^®^. Luminescence was measured in a 2104 EnVision^®^ multilabel plate reader (PerkinElmer). The activity of test compounds was normalized to controls from the same plate. Dose–response curves and CC_50_ values, *i.e*., the concentration of compound required to inhibit cell growth by 50%, were calculated using GraphPad Prism version 9.4.1 for MAC OS (GraphPad Software). Assays were performed as one to three experimental replicates, each as a singleton.

#### Time-kill Assays

This was assessed using the CellTiter-Glo^®^ reagent (Promega, Cat. No. G7572) to measure levels of ATP as a real-time indicator of parasite viability.^48^ Test compounds were serially diluted in DMSO across eight concentrations ranging from 4 μM to 0.87 nM. Compounds were added to the parasite cultures as solutions in DMSO. Trypanosomes in log-phase growth were suspended at 2×10^5^ parasites/ml in HMI-9 medium under continuous agitation. Then, cells were dispensed into 96-well plates (Corning^®^, Cat. No. 3903) (50 μL/well) containing 1 μl of test compound in 1% DMSO and 50 μl/well of fresh medium. Assay plates were incubated at 37 °C and 5% CO_2_ and at each selected time, 50 μl/well of CellTiter-Glo^®^ reagent was added. Plates were gently shaken and incubated in the dark for 10 min. Luminescence was measured in a 2104 EnVision^®^ multilabel plate reader (PerkinElmer). The activity of test compounds was normalized to DMSO controls from the same plate. Dose– response curves and EC_50_ values were calculated using GraphPad Prism version 9.4.1 for MAC OS (GraphPad Software). Assays were performed as three experimental replicates, each as a duplicate.

### Measurement of AcTub and Total α-Tub Levels in HEK-293 Cells

These assays have been described previously.^30, 33^

### *In Vivo* Efficacy Experiments

*Ethics Statement.* Experiments were carried out in accordance with protocols approved by the Institutional Animal Care and Use Committee (IACUC) at UCSD. Approval from UCSD-IACUC was granted under the Animal Welfare Act and Regulations (AWAR) Act and the United States Public Health Service Policy on Humane Care and Use of Laboratory Animals.

#### Mouse Model of T. brucei Infection

Six-week-old wild-type BALB/c female mice were infected with 1×10^5^ *T. b. brucei* Lister 427 cells by i.p. injection. Mice were divided into groups of five, and 48 h after infection, compounds were administered i.p. in 9% DMSO and 91% corn oil at the doses indicated in the Results section. Pentamidine (Sigma-Aldrich, Cat No. P0547) was used as reference drug. Infected control mice groups were those that received either no treatment or given vehicle only (vehicle controls).

Blood parasitemia was monitored by microscopic examination of tail blood every 24 to 48 h starting at 48 h post-infection and up to 16 days post-infection. Specifically, 5 µl blood was withdrawn and diluted immediately with 5 µl heparin solution (0.5 mg/ml in sterile water). Samples were diluted a further 100-fold in PBS at 37 °C and the parasites counted using a Neubauer chamber. The lower limit for accurate counting of cells in a Neubauer chamber is 2.5×10^5^ cells/ml. Mice were monitored every day for weight loss, general appearance and behaviour. Mice showing impaired health status and/or distress, such as reduced grooming and socialization, were humanely euthanized.

#### Tolerability Studies

These were conducted in 8-week-old BALB/c mice and entailed a preliminary assessment in one mouse followed by a more rigorous study in groups comprising six mice of each sex. Mice were monitored twice daily for 48 h post-administration for overt signs of intolerance such as altered grooming and socialization behaviors. Upon the discovery of either of these signs, the affected mice were humanely euthanized.

### Determination of Plasma and Brain Drug Concentrations

PK analysis of compound **4** was conducted under contract by Touchstone Biosciences (Plymouth Meeting, PA). Briefly, male CD-1 mice (n=3 per time point) received 2 mg/kg i.p. of **4** dissolved in a 10% DMSO:40% polyethylene glycol:50% H_2_O solution. Groups of mice were sacrificed at 0.5, 1, 2, 4 and 8 h after compound dosing, and blood was collected from each animal into Greiner MiniCollect K_2_EDTA tubes on ice and then centrifuged at 15,000 *g* for 5 min to obtain plasma samples. All plasma samples were stored at –70 °C until analysis. The whole brain was harvested at each time point and was rinsed in water followed by homogenization in a tissue homogenizer in PBS buffer, pH 7.4 (3:1 volume/brain weight). The brain homogenate samples were stored at –70°C until analysis. Compound levels in plasma and brain were determined by LC-MS/MS using protocols previously described.^25, 49^

## Supporting Information

**Figure S1.** Phylogenetic comparison of α- and β-tubulins from key trypanosomatid species compared to humans.

**Figure S2.** Time-kill assays for *T. b. brucei* Lister 427 in the presence of the TPDs **3** and **4**, and pentamidine.

**Figure S3.** Validation of the homology model of *T. brucei* MT with ERRAT, VERIFY and PROVE using SAVES v6.0.

**Table S1.** Complete list of 1,2,4-Triazolo[1,5-a]pyrimidine analogues **1** – **123** used in the study.

**Table S2.** Parasitemia levels (expressed as number of trypanosomes/ml) of BALB/c mice infected with *T. brucei* and either not treated or treated with vehicle only (9% DMSO and 91% corn oil).

**Table S3.** Parasitemia levels of BALB/c mice infected with T. brucei and treated with pentamidine at 4 mg/kg on day 2 post-infection.

**Table S4.** Parasitemia levels of BALB/c mice infected with *T. brucei* and treated with TPD **3** at 5, 7.5, and 10 mg/kg on day 2 post-infection.

**Table S5.** Parasitemia levels of BALB/c mice infected with T. brucei and treated with TPD **4** at 5, 7.5, and 10 mg/kg on day 2 post-infection.

## Acknowledgments

Financial support for this work has been provided by the NIH-funded research projects R21AI133394 and R21AI141210.

## Abbreviations

HAT: Human African trypanosomiasis
MT: Microtubules
TPD: triazolopyrimidine
SAR: structure-activity relationship
PK: pharmacokinetics
AcTub: acetylated α-tubulin
i.p.: intraperitoneal.

## Table of Contents graphic

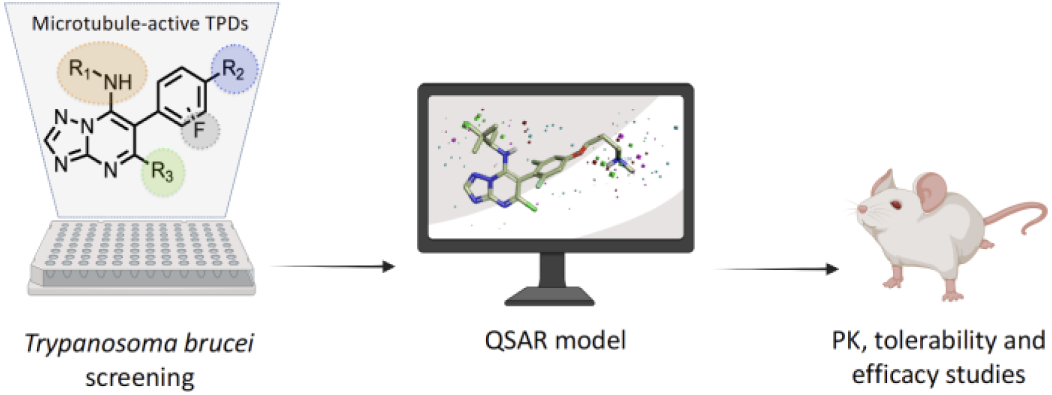

