## Supplemental Materials for "Structure-Activity Relationships, Tolerability and Efficacy of Microtubule-Active 1,2,4-Triazolo[1,5-*a*]pyrimidines as Potential Candidates to Treat Human African Trypanosomiasis"

**Table of contents**

**Figure S1.** Phylogenetic comparison of α- and β-tubulins from key trypanosomatid species compared to humans.

**Figure S2**. Time-kill assays for *T. b. brucei* Lister 427 in the presence of the TPDs **3** and **4**, and pentamidine.

**Figure S3.** Validation of the Homology model of *T. brucei* MT with ERRAT, VERIFY and PROVE using SAVES v6.0.

**Table S1.** Complete list of 1,2,4-Triazolo[1,5-*a*]pyrimidine analogues **1** – **123** used in the study. (Table S1 is included in the Supporting Information as a separate .csv file)

**Table S2.** Parasitemia levels (expressed as number of trypanosomes/ml) of BALB/c mice infected with *T. brucei* and either not treated or treated with vehicle only (9% DMSO and 91% corn oil).

**Table S3.** Parasitemia levels of BALB/c mice infected with *T. brucei* and treated with pentamidine at 4 mg/kg on day 2 post-infection.

**Table S4.** Parasitemia levels of BALB/c mice infected with *T. brucei* and treated with TPD **3** at 5, 7.5, and 10 mg/kg on day 2 post-infection.

**Table S5.** Parasitemia levels of BALB/c mice infected with *T. brucei* and treated with TPD **4** at 5, 7.5, and 10 mg/kg on day 2 post-infection.

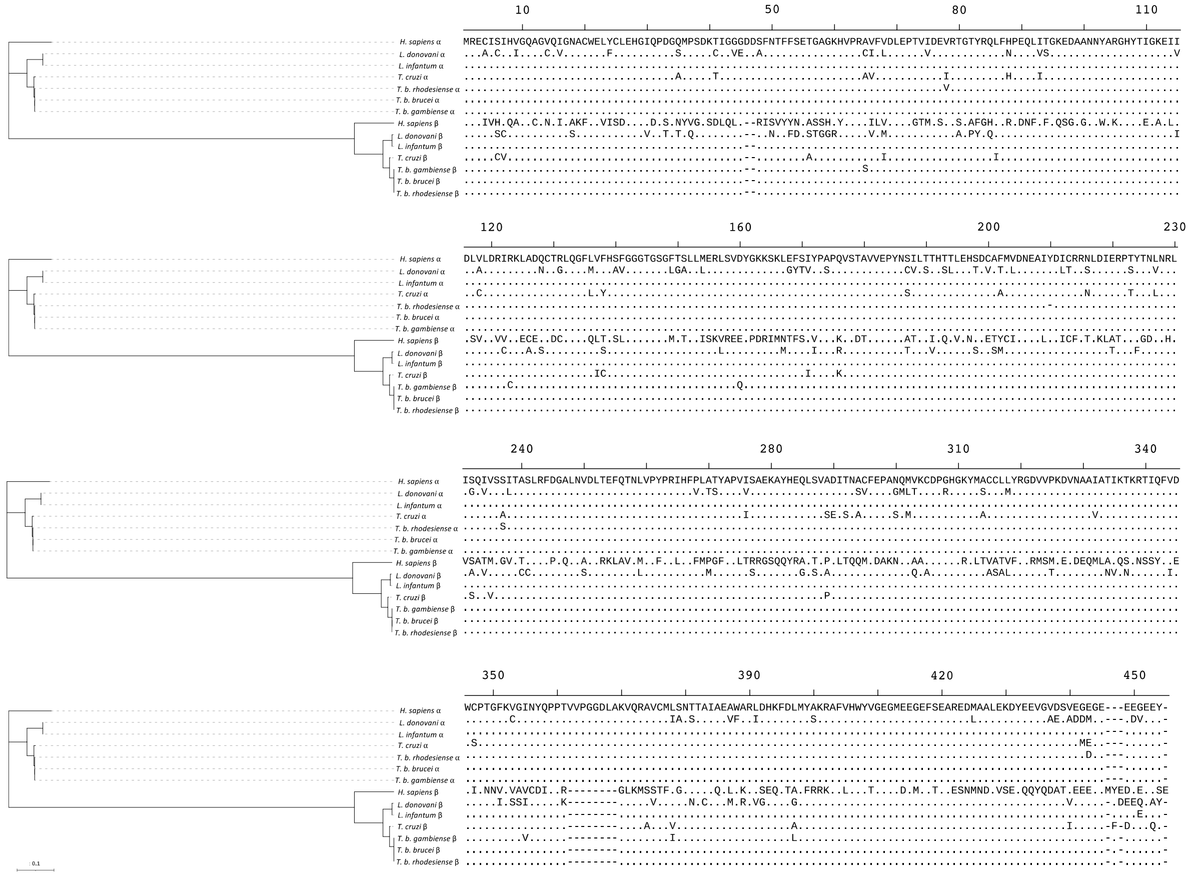

**Figure S1. Phylogenetic comparison of α- and β-tubulins from key trypanosomatid compared to humans.** Full-length alignment of α- and β-tubulin amino acid sequences from *Trypanosoma brucei brucei*, *Trypanosoma brucei gambiense*, *Trypanosoma brucei*

*rhodesiense*, *Trypanosoma cruzi*, *Leishmania donovani*, *Leishmania infantum* and *Homo sapiens*. The alignment was built using MUSCLE[1, 2] and served as the input for the construction of a maximum likelihood phylogenetic tree using IQ-TREE2[3]. The interactive tree of life program (iTOL)[4] was used to visualise the phylogenetic tree and sequence alignment. The scale bar represents an approximate value for the amino acid substitution rate.

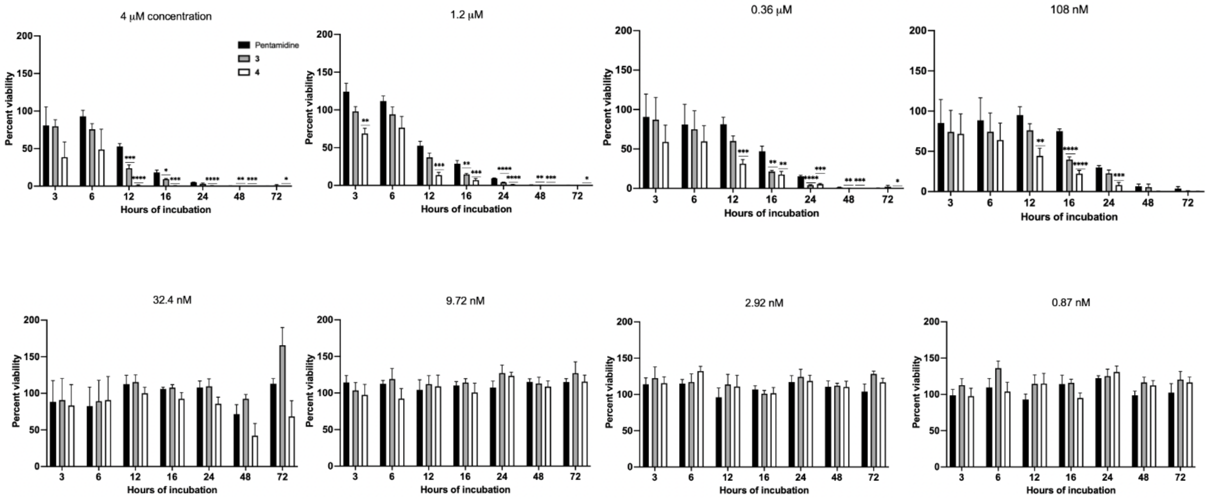


**Figure S2. Time-kill assays for *T. b. brucei* Lister 427 in the presence of the TPDs 3 and 4, and pentamidine.** *T. brucei* was exposed to 0.87 nM through 4 µM compound as a function of time (3 – 72 h). All assays were performed as three experimental replicates, each in duplicate. The activity of test compounds was normalized to DMSO controls from the same plate and means ± SEM values are shown. The Student's unpaired t-test was used to compare the data for the TPDs vs. those for pentamidine: *$p < 0.05$, **$p < 0.01$, ***$p < 0.001$, ****$p < 0.0001$.

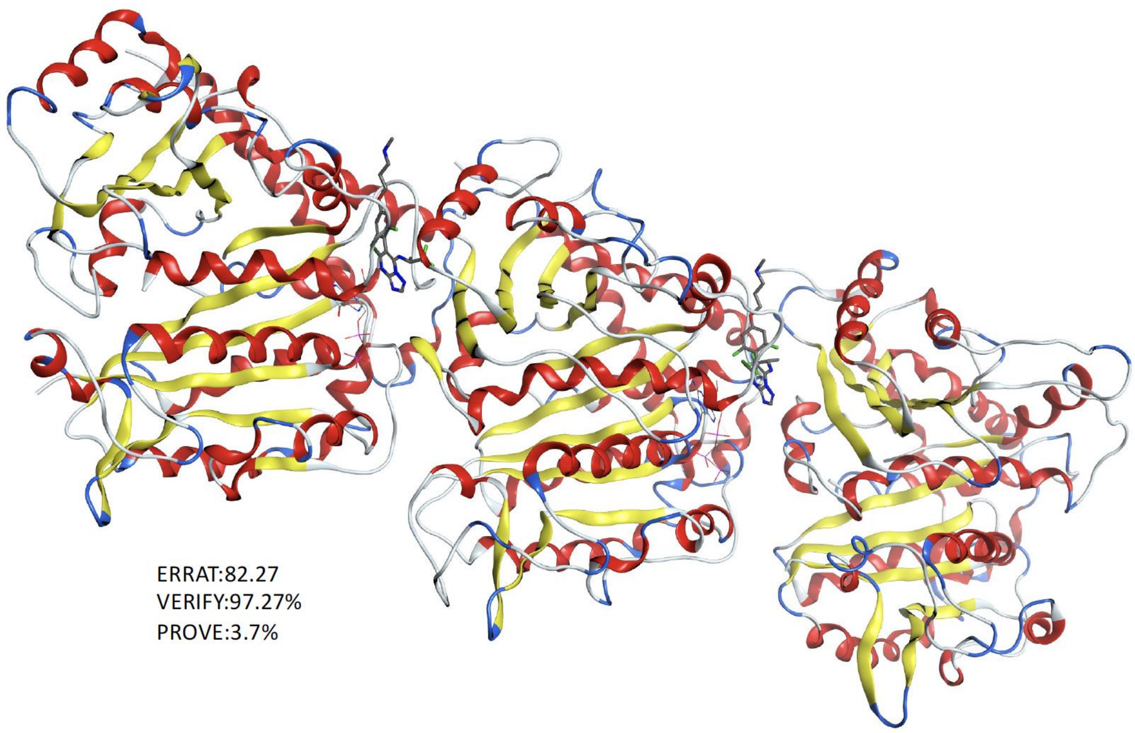


**Figure S3.** Validation of the Homology model of *T. brucei* MT with ERRAT, VERIFY and PROVE using SAVES v6.0.

**Table S2.** Parasitemia levels (expressed as number of trypanosomes/ml) of BALB/c mice infected with *T. brucei* and either not treated or treated with vehicle only (9% DMSO and 91% corn oil).

| **Group** | **Untreated** | | | | | **Vehicle** | | | | |
|---|---|---|---|---|---|---|---|---|---|---|
| Mouse / Parasites/ml | 1 | 2 | 3 | 4 | 5 | 1 | 2 | 3 | 4 | 5 |
| Day 2 | n.d. | n.d. | n.d. | n.d. | n.d. | n.d. | n.d. | n.d. | n.d. | n.d. |
| Treatment (μL) | - | - | - | - | - | 100 | 100 | 100 | 100 | 100 |
| Day 3 | $6.72x10^8$ | $6.42x10^8$ | $4.28x10^8$ | $5.76x10^8$ | $3.80x10^8$ | $7.2x10^7$ | $6.32x10^8$ | $4.20x10^8$ | $4.56x10^8$ | $4.44x10^8$ |

**Table S3.** Parasitemia levels of BALB/c mice infected with *T. brucei* and treated with pentamidine at 4 mg/kg on day 2 post-infection.

| **Group** | **Vehicle control** | | | | | **Pentamidine** | | | | |
|---|---|---|---|---|---|---|---|---|---|---|
| Mouse / Parasites/ml | 1 | 2 | 3 | 4 | 5 | 1 | 2 | 3 | 4 | 5 |
| Day 2 | $1.4x10^7$ | NA | $1.8x10^7$ | $1.0x10^7$ | $8x10^6$ | $6x10^6$ | $4x10^6$ | $2x10^6$ | $2x10^6$ | $4x10^6$ |

| | | | | | | | | | | |
|---|---|---|---|---|---|---|---|---|---|---|
| Dose (mg/kg) on day 2 | - | - | - | - | - | 4 | 4 | 4 | 4 | 4 |
| Day 3 | 6.88x$10^{8}$ | 6.62x$10^{8}$ | 6.66x$10^{8}$ | 4.54x$10^{8}$ | 5.84x$10^{8}$ | NA | NA | NA | NA | NA |
| Day 14 | - | - | - | - | - | NA | n.d. | n.d. | n.d. | n.d. |
| Day 15 | - | - | - | - | - | n.d. | n.d. | n.d. | n.d. | n.d. |

**Table S4.** Parasitemia levels of BALB/c mice infected with *T. brucei* and treated with TPD **3** at 5, 7.5, and 10 mg/kg on day 2 post-infection.

| **Group** | **Vehicle control** | | | | | **5 mg/kg** | | | | | **7.5 mg/kg** | | | | | **10 mg/kg** | | | | |
|---|---|---|---|---|---|---|---|---|---|---|---|---|---|---|---|---|---|---|---|---|
| Mouse / Parasites/ml | 1 | 2 | 3 | 4 | 5 | 1 | 2 | 3 | 4 | 5 | 1 | 2 | 3 | 4 | 5 | 1 | 2 | 3 | 4 | 5 |
| Day 2 | 3.7x$10^{7}$ | NA | 4.7x$10^{7}$ | 5.1x$10^{7}$ | 5.7x$10^{7}$ | 3.0x$10^{7}$ | 5.6x$10^{7}$ | 3.7x$10^{7}$ | 3.9x$10^{7}$ | 2.7x$10^{7}$ | 1x$10^{7}$ | 1x$10^{6}$ | 2.4x$10^{7}$ | 1.4x$10^{7}$ | 1.3x$10^{7}$ | 2.5x$10^{7}$ | 4.0x$10^{7}$ | 3.5x$10^{7}$ | 2.6x$10^{7}$ | 3.4x$10^{7}$ |

| Dose (mg/kg) on day 2 | - | - | - | - | - | 5 | 5 | 5 | 5 | 5 | 7.5 | 7.5 | 7.5 | 7.5 | 7.5 | 10 | 10 | 10 | 10 | 10 |
|---|---|---|---|---|---|---|---|---|---|---|---|---|---|---|---|---|---|---|---|---|
| Day 3 | 7.23 x$10^8$ | NA | 1x$10^9$ | 8.87 x$10^8$ | 6.87 x$10^8$ | 3x$10^6$ | 1x$10^6$ | 1.4x $10^7$ | 3.4x $10^6$ | 3x$10^6$ | NA | NA | NA | NA | NA | NA | NA | NA | NA | NA |
| Day 4 | - | 7.2x $10^7$ | - | - | - | 7.1x $10^7$ | 2.4x $10^7$ | 1.71 x$10^8$ | 6.7x $10^7$ | 3.4x $10^7$ | NA | NA | NA | NA | NA | NA | NA | NA | NA | NA |
| Day 5 | - | 9.81 $10^8$ | - | - | - | 2.01 x$10^8$ | 1.77 x$10^8$ | 1.03 x$10^9$ | 7.12 x$10^8$ | 4.21 x$10^8$ | NA | NA | NA | NA | NA | NA | NA | NA | NA | NA |
| Day 6 | - | - | - | - | - | n.d. | n.d. | - | - | n.d. | NA | NA | NA | NA | NA | NA | NA | NA | NA | NA |
| Day 7 | - | - | - | - | - | - | - | - | - | - | NA | NA | NA | NA | NA | NA | NA | NA | NA | NA |
| Day 8 | - | - | - | - | - | - | - | - | - | - | 1.6x $10^7$ | NA | 5x$10^6$ | NA | NA | NA | NA | 6.6x $10^5$ | NA | NA |
| Day 9 | - | - | - | - | - | - | - | - | - | - | 5x$10^7$ | NA | 3.6x $10^7$ | NA | 3x$10^6$ | NA | NA | 1.47 x$10^8$ | 4.69 x$10^8$ | NA |

| | | | | | | | | | | | | | | | | | | | | |
|---|---|---|---|---|---|---|---|---|---|---|---|---|---|---|---|---|---|---|---|---|
| 2nd Dose (mg/gg) on day 9 | - | - | - | - | - | - | - | - | - | - | 7 | 7 | 7 | 7 | 7 | 5 | 5 | 5 | 5 | 5 |
| Day 10 | - | - | - | - | - | - | - | - | - | - | NA | NA | - | NA | - | - | NA | - | - | NA |
| Day 11 | - | - | - | - | - | - | - | - | - | - | n.d. | n.d. | - | n.d. | - | - | NA | - | - | NA |
| Day 12 | - | - | - | - | - | - | - | - | - | - | n.d. | n.d. | - | n.d. | - | - | n.d. | - | - | n.d. |
| Day 13 | - | - | - | - | - | - | - | - | - | - | $8x10^{6}$ | $5x10^{6}$ | - | $3x10^{6}$ | - | - | $1.5x10^{6}$ | - | - | $6.6x10^{7}$ |
| Day 14 | - | - | - | - | - | - | - | - | - | - | $1.18x10^{8}$ | $8x10^{6}$ | - | $9x10^{6}$ | - | - | n.d. | - | - | n.d. |
| Day 15 | - | - | - | - | - | - | - | - | - | - | $4.76x10^{8}$ | $1.94x10^{8}$ | - | $9.4x10^{7}$ | - | - | $2.69\ 10^{8}$ | - | - | $5.91\ 10^{8}$ |

**Table S5.** Parasitemia levels of BALB/c mice infected with *T. brucei* and treated with TPD **4** at 5, 7.5, and 10 mg/kg on day 2 post-infection.

| **Group** | **Vehicle control** | **5 mg/kg** | **7.5 mg/kg** | **10 mg/kg** |
|---|---|---|---|---|

| Mouse / Parasites/ml | 1 | 2 | 3 | 4 | 5 | 1 | 2 | 3 | 4 | 5 | 1 | 2 | 3 | 4 | 5 | 1 | 2 | 3 | 4 | 5 |
|---|---|---|---|---|---|---|---|---|---|---|---|---|---|---|---|---|---|---|---|---|
| Day 2 | 6x10$^{6}$ | 1.5x10$^{7}$ | 1.4x10$^{7}$ | 2.3x10$^{7}$ | 3.9x10$^{7}$ | 2.7x10$^{7}$ | 1.8x10$^{7}$ | 2.1x10$^{7}$ | 1.3x10$^{7}$ | 1.6x10$^{7}$ | 2.7x10$^{7}$ | 3.1x10$^{7}$ | 1.0x10$^{7}$ | 2.9x10$^{7}$ | 1.8x10$^{7}$ | 3.6x10$^{7}$ | 1.7x10$^{7}$ | 1.7x10$^{7}$ | 3.6x10$^{7}$ | 3.0x10$^{7}$ |
| Dose (mg/kg) on day 2 | - | - | - | - | - | 5 | 5 | 5 | 5 | 5 | 7.5 | 7.5 | 7.5 | 7.5 | 7.5 | 10 | 10 | 10 | 10 | 10 |
| Day 3 | 5.14x10$^{8}$ | 6.82x10$^{8}$ | 3.16x10$^{8}$ | 3.80x10$^{8}$ | 2.46x10$^{8}$ | 1.6x10$^{7}$ | 1x10$^{7}$ | 6.4x10$^{7}$ | 2.6x10$^{7}$ | 4x10$^{6}$ | 1.34x10$^{6}$ | 6.6x10$^{5}$ | 6.6x10$^{5}$ | 2.66x10$^{6}$ | 2.66x10$^{6}$ | NA | NA | NA | NA | NA |
| Day 4 | - | - | - | - | - | 8.6x10$^{7}$ | 2.44x10$^{8}$ | 6.52x10$^{8}$ | 2.78x10$^{8}$ | 4.2x10$^{7}$ | 7.6x10$^{7}$ | 4x10$^{6}$ | NA | 4.2x10$^{7}$ | 2.2x10$^{7}$ | NA | NA | NA | NA | NA |
| Day 5 | - | - | - | - | - | - | - | - | - | - | - | - | - | - | - | n.d. | n.d. | n.d. | n.d. | n.d. |
| Day 6 | - | - | - | - | - | - | - | - | - | - | - | - | - | - | - | NA | NA | NA | NA | NA |

| | | | | | | | | | | | | | | | | | | | | |
|---|---|---|---|---|---|---|---|---|---|---|---|---|---|---|---|---|---|---|---|---|
| Day7 | - | - | - | - | - | - | - | - | - | - | - | - | - | - | - | n.d. | n.d. | n.d. | n.d. | n.d. |
| Day 8 | - | - | - | - | - | - | - | - | - | - | - | - | - | - | - | 6.6x10$^{5}$ | 2x10$^{6}$ | NA | 6.6x10$^{5}$ | 6.6x10$^{5}$ |
| Day 9 | - | - | - | - | - | - | - | - | - | - | - | - | - | - | - | 6x10$^{6}$ | 1.2x10$^{7}$ | 1.6x10$^{7}$ | 2x10$^{6}$ | 1.1x10$^{7}$ |
| Dose (mg/kg) on day 9 | - | - | - | - | - | - | - | - | - | - | - | - | - | - | - | 10 | 10 | 10 | 10 | 10 |
| Day 10 | - | - | - | - | - | - | - | - | - | - | - | - | - | - | - | NA | NA | NA | NA | NA |
| Day 11 | - | - | - | - | - | - | - | - | - | - | - | - | - | - | - | n.d. | n.d. | n.d. | n.d. | n.d. |
| Day 12 | - | - | - | - | - | - | - | - | - | - | - | - | - | - | - | NA | 2x10$^{6}$ | NA | NA | NA |
| Day 13 | - | - | - | - | - | - | - | - | - | - | - | - | - | - | - | n.d. | n.d. | n.d. | n.d. | n.d. |

| | | | | | | | | | | | | | | | | | | | | |
|---|---|---|---|---|---|---|---|---|---|---|---|---|---|---|---|---|---|---|---|---|
| Day 14 | - | - | - | - | - | - | - | - | - | - | - | - | - | - | - | NA | NA | 2x10$^{6}$ | NA | NA |
| Day 15 | - | - | - | - | - | - | - | - | - | - | - | - | - | - | - | n.d. | n.d. | n.d. | n.d. | n.d. |
| Day 16 | - | - | - | - | - | - | - | - | - | - | - | - | - | - | - | 6.6 x10$^{5}$ | 5x10$^{6}$ | 1.3 x10$^{7}$ | NA | 6.6 x10$^{5}$ |

NA = Not applicable due to parasitemia levels being below the limit of detection (*i.e.*, 2.5×10$^{5}$ trypanosomes/ml).
n.d. = not determined
